# Spatial multi-omics implicates vascular adipocytes proximal to tertiary lymphoid structures in human aortic aneurysm and B-cell autoreactivity

**DOI:** 10.64898/2026.10.13.758932

**Authors:** Akshaya K. Meher, Weihua Huang, Dean J. Yamaguchi, Changhong Yin, Madison M. Swyers, Gabriel Dubis, Li V. Yang, Mark D. Mannie, Coleen A. McNamara

## Abstract

**Objective:** Perivascular Adipose Tissue (PVAT), tertiary lymphoid structures (TLS), and the B lymphocytes they harbor have been implicated in regulating vascular diseases, including abdominal aortic aneurysm (AAA). Yet, the relationship of adipose tissue to TLS and B-cell responses is poorly understood.

**Approach and Results:** Here, using multi-omics approaches, we characterized the potential interplay of vascular adipocytes and TLSs in shaping microenvironments conducive to pathogenic B-cell immunity in human AAA pathology. Single-nucleus RNA sequencing identified white adipocytes expressing transcripts for lipogenesis, lipolysis, and extracellular remodeling. Unexpectedly, these adipocytes expressed haptotactic and chemotactic factors such as thrombospondin 1 and autotaxin, and extracellular matrix proteins with potential to recruit CD44+ leukocytes within AAAs. Immunohistology revealed two anatomical compartments for the adipocytes within AAAs: PVAT and the medial-adventitial region. Medial-adventitial adipocytes were frequently associated with CD44+ TLSs and were largely absent from control non-aneurysmal abdominal aortas. Visium HD spatial transcriptomics revealed mature TLSs recapitulated the organization of functional secondary lymphoid structures, with multiple B-cell follicles clustered to form a nodular structure partly covered by a layer of smooth muscle cells. The TLSs harbored isotype-switched B cells expressing IgG1, IgG3, or IgA, with dominant kappa chain usage and IgG antibodies to the aortic wall. Myeloid cells in AAA were implicated in promoting B-cell effects, as they were the primary producers of B-cell activating factor (BAFF), a cytokine that promotes B-cell infiltration, survival, and function.

**Conclusion:** Altogether, our results support a model in which new adipocytes in the medial-adventitial layer elicit inflammatory pathways that promote early TLS formation, and BAFF produced by myeloid cells promotes the survival and maturation of vascular wall-antigen-exposed B cells, leading to mature TLS formation, production of antibodies to AAA antigens, fueling inflammation and AAA growth.

**Graphical Abstract:** 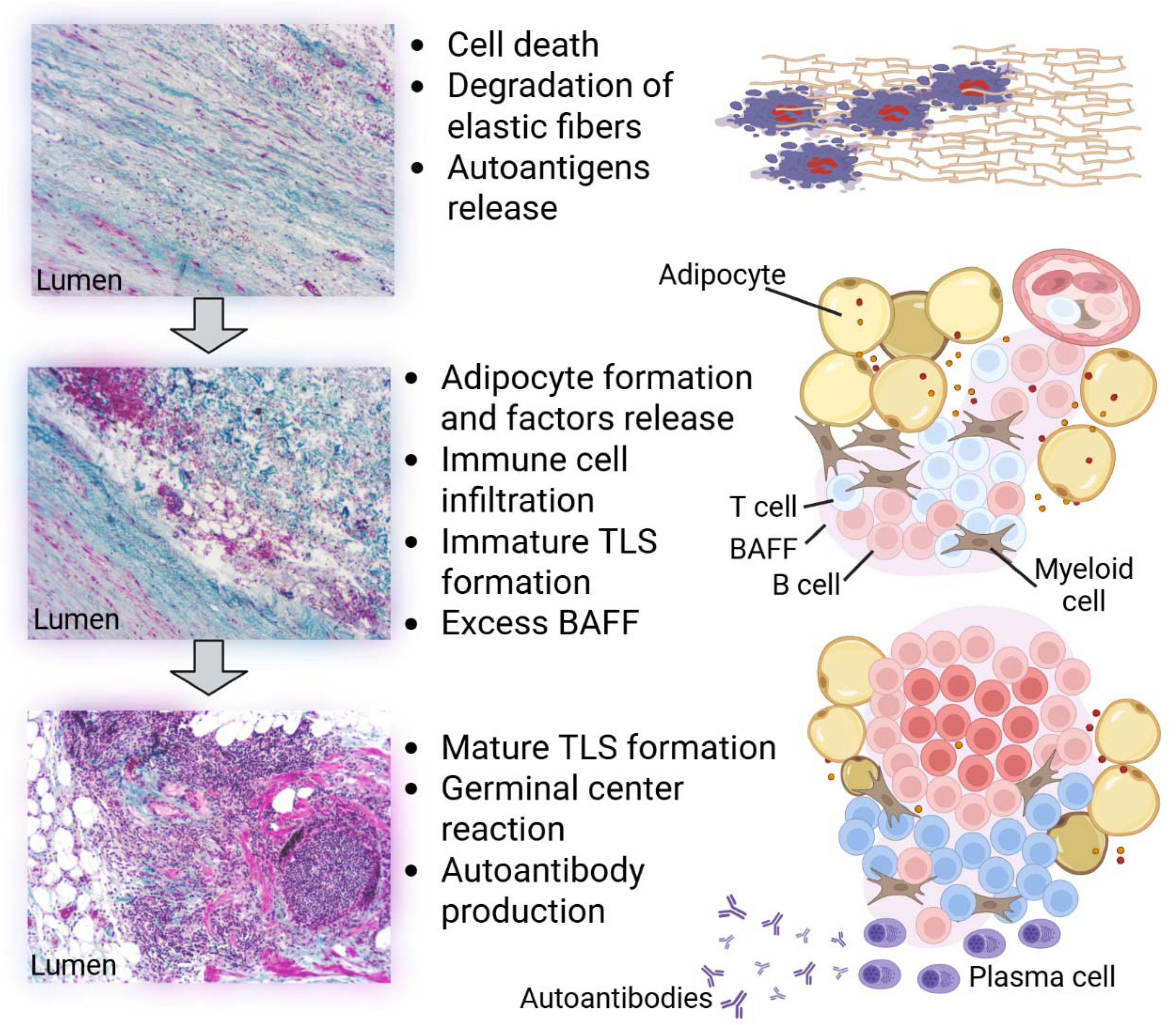

**Clinical Implications:** Despite multiple drugs showing efficacy in preclinical animal models, no non-surgical treatments are available for AAAs. Therefore, we re-evaluated human aortic aneurysm pathology using new technologies to better understand human aneurysm pathogenesis. The results revealed formation of novel adipocytes in the medial-adventitial region of human AAAs. These vascular adipocytes exacerbate inflammation by attracting an arsenal of immune cells to the medial-adventitial region. B-cell activating factor (BAFF) further exacerbates the B-cell immune response by supporting the formation of mature tertiary lymphoid structures (TLSs) and production of antibodies against AAA tissue. While more studies are needed to confirm these findings, this work opens new avenues of investigation for potential therapeutics for treating small AAAs. Particularly, repurposing anti-adipogenesis drugs and anti-BAFF therapeutics may attenuate B-cell immune responses, inflammation, and AAA growth.

## Introduction

Aortic aneurysm (AA) is defined as a ≥50% enlargement of the aorta^1^, and is a significant life-threatening risk factor for elderly males, as aneurysm rupture causes death in 80-90% of cases^1,2^. Abdominal AAs (AAAs) are the most prevalent form of AA^3^. Yet, to date, there are no non-surgical treatments available for AAAs despite multiple drugs demonstrating efficacy in preclinical animal models. Murine models of AAAs have advanced our understanding of AAA formation and growth, but these models have several limitations^4,5^ and understanding the pathogenesis of human aneurysms is key to developing effective therapies. Therefore, re-evaluating human aortic aneurysm pathology using new technologies is crucial for developing new models and treatment options.

A wealth of evidence supports the role of inflammatory changes in vascular intramural cells, such as smooth muscle cells, endothelial cells, and fibroblasts, in aneurysm growth in both human and murine experimental AAAs, contributing to AAA pathogenesis. Notably, both inflammatory phenotypes and aneurysm lesion formation are attenuated in murine hosts that lack T-cells^6^ or B-cells^7^, suggesting a critical role of lymphocytes in driving inflammatory activities of aortic mural cells and ultimately in eliciting AAA lesion formation. Previous work has shown that 1) B cells are present in the human and in murine AAAs; 2) pharmacological depletion of B cells protects mice from AAA; 3) pharmacological depletion of the TNF cytokine B-cell activating factor (BAFF), a critical cytokine for differentiation of naïve B-2 cells to mature B-2 cells, attenuates the growth of existing AAAs in a murine model; 4) BAFF and its receptors are expressed in human tertiary lymphoid structures (TLSs) found in human AAAs.

The appearance of TLSs in non-lymphoid organs is a hallmark of autoimmunity^8^. TLSs are organized aggregates of B cells and T cells, formed after chronic inflammation and generate tissue-specific autoantibodies that exacerbate inflammation^9,10^. Studies with BAFF deficiency or sufficiency revealed that, at the site of chronic inflammation, infiltrated naïve autoreactive anergic B cells, after exposure to excess autoantigens, survive with the help of BAFF, form germinal centers, and differentiate into plasmablasts, followed by autoantibody-secreting plasma cells^11^. While the presence of TLSs in AAAs has been reported before^12,13^, the ontogeny of TLS in AAAs is not clearly understood. Furthermore, it is unclear which vascular mural cells promote lymphocyte organization into TLSs, what is the source of BAFF, and how BAFF supports TLSs in human AAAs.

To understand the key players in TLS formation in AAA, we utilized a targeted multi-omic approach including single-nucleus (sn) RNA-seq, Visium HD spatial transcriptomics, and NanoString spatial proteomics, along with analysis of an existing single-cell (sc) RNA-seq dataset from human AAA and control biospecimens. We discovered that some adipocytes function as non-canonical TLS organizers by synthesizing haptotactic and chemotactic factors such as thrombospondin 1, autotaxin, and extracellular matrix proteins that attract and promote CD44+ leukocyte cluster formation. Additionally, BAFF supports TLS maturation and B-cell generation of autoantibodies targeting the aorta and damaged tissues in human AAAs.

## MATERIALS AND METHODS

### Data availability

SnRNA-seq data from 4 AAA patients were deposited in the Gene Expression Omnibus (GEO) repository with accession number GSE350968. Some analyzed data are provided in the supplemental tables, and other data are available upon request.

### Human aortic tissue acquisition

Discarded AAA biospecimens from patients undergoing open repair at East Carolina University Health, NC, USA were collected under an approved Institutional Review Board CIRB 20-001312. Written informed consents were obtained from AAA patients before tissue collection. Following project approval, non-aneurysmal control abdominal aortas from organ donors were obtained from the National Disease Research Interchange, PA, USA. For both AAA and control aortas, adults (males and females) aged 50 to 88, irrespective of race, were selected. While AAA biospecimens were available within 2-3 hours of removal, control abdominal aortas were shipped in ice-cold Dulbecco’s Modified Eagle Medium (DMEM) and were available within 24 hours of removal. Once available, the biospecimens were washed with cold Dulbecco’s Phosphate-Buffered Saline (DPBS) and cut into 3 pieces under sterile conditions: one part was frozen and stored at −80 °C, another part was fixed for 48 hours in Buffered Zinc Formalin (# 22-050-259, Thermo Scientific), and the remaining part was used for *ex vivo* culture.

### Single-nucleus RNA sequencing (snRNA-seq)

Initially, snRNA-seq studies on one 100 mg AAA frozen specimen were performed by Novogene, USA. Subsequently, snRNA-seq studies on AAAs from four patients (including the one that was performed by Novogene) were performed at East Carolina University. Briefly, two 100 mg frozen tissue samples were cut from different sites on the frozen AAA specimen, and nuclei were prepared in two separate tubes using 10x Genomics’s Chromium Nuclei Isolation Kit with RNase Inhibitor. Next, nuclei preparations from the two tubes were pooled together. Single nuclei prepared were appraised using the Countess 3 FL (ThermoFisher) for both quantity and quality checks and subjected to library construction using the Chromium Single Cell 3’ Gene Expression kit and the Chromium Controller (10x Genomics), according to the manufacturer’s protocol. Sequencing of the pooled libraries was performed on the Illumina NextSeq 2000 platform. Raw sequencing reads in FASTQ files were obtained using cellranger (v10.0.0, 10x Genomics) mkfastq function. Here, we included the raw data obtained from one AAA specimen from Novogene. Thus, we had 5 AAAs from 4 patients.

The alignment to the human GRCh38 reference genome and the subsequent quantification were conducted using the cellranger count function. Subsequently, Seurat (v5.5.0, https://github.com/satijalab/seurat) package was employed in RStudio (Build 526) with R (v4.6.0) for bioinformatics data analysis. Briefly, single nuclei were first filtered with nFeature_RNA (more than 300 but less than 6000) and percentage of mitochondrial reads (percent.mt, less than 15%). Doublets or multiplets were eliminated using DoubletFinder (v2.0.4, https://github.com/chris-mcginnis-ucsf/DoubletFinder); and batch effects were corrected using Harmony (v2.0.5, https://github.com/immunogenomics/harmony). Gene expression data were then aggregated, normalized, and variance-stabilized with sctransform-regularized negative binomial regression (SCTransform function in Seurat), regressing out mitochondrial reads. Dimensionality reduction was performed by principal component analysis (PCA) and uniform manifold approximation and projection (UMAP) embedding (dims = 1:30) with RunPCA and RunUMAP, respectively. Cell clusters were subsequently identified with FindNeighbors (dims = 1:30) and FindClusters (resolution = 0.3); and annotated based on expression of cell type-specific gene biomarkers. Differentially expressed biomarkers among cell types were assessed with FindAllMarkers or FindMarkers for genes detected in at least 10% of cells using a log(foldchange) threshold at 0.25. Bonferroni-adjusted p-values were used to determine significance at a false discovery rate (FDR) < 0.05. The clustered cell types were manually annotated with their differentially expressed gene markers and their related Kyoto Encyclopedia of Genes and Genomes (KEGG) pathway analysis (adjusted p-value <0.05) using the National Institutes of Health’s DAVID Bioinformatics (https://davidbioinformatics.nih.gov/). CellChat (v2.2.0.9001, doi: 10.1038/s41596-024-01045-4) was finally used for ligand-receptor inferred analysis of cell-cell communication. For the remaining analyses, we used default parameters.

### 10x Genomics Visium HD spatial transcriptomics

Sections (10□µm) of Formalin-Fixed Paraffin-Embedded (FFPE) tissue blocks were placed on standard glass slides, stained with Hematoxylin and Eosin (H&E), and imaged under MoticEasyScan Pro system (Motic), following the 10x Genomics protocol: Visium CytAssist Spatial Gene Expression for FFPE. Samples were then transferred to a Visium HD slide with CytAssist and processed for library preparation using the Visium HD Spatial Gene Expression Reagent, following the user guide (10x Genomics). Sequencing was performed on the Illumina NextSeq 2000 platform, with raw sequencing reads in FASTQ files generated by the on-instrument DRAGEN (v3.8.4). The gene expression quantification was obtained by using the human GRCh38 reference transcriptome and spaceranger (v4.0.1, 10x Genomics) count function. Seurat was employed for the subsequent bioinformatics data analysis, similar to the scRNAseq above. By aggregating Visium HD 2×2-µm bins, we explored 8×8 and 16×16-µm binned data to balance capture noise and cellular resolution. The spatial gene expression data were deconvolved with the Robust Cell Type Decomposition (RCTD, https://doi.org/10.1038/s41587-021-00830-w) algorithm, using clustered and annotated cell types in our scRNA-seq as reference and the spacexr package (https://github.com/dmcable/spacexr).

### NanoString spatial proteomics

Five-micrometer cross-sections of formalin-fixed paraffin-embedded human control aorta with cellular infiltrates in the adventitia and AAA with TLSs were obtained from OriGene, MD, USA. The cross-sections were sent to NanoString Technologies, WA, USA, for spatial proteomics studies using GeoMx Immuno-Oncology Protein Panels, developed by NanoString/Bruker Spatial Biology. This panel has UV-cleavable DSP (Digital Spatial Profiler)-tagged antibodies for 79 distinct targets including 3 housekeeping proteins (GAPDH, Histone H3 & Ribosomal protein S6) and 3 negative control Isotypes (Mouse IgG1, Mouse IgG2a & Rabbit IgG). After the assay, three normalizations: area normalization, Housekeeper normalization, and signal-to-noise normalization were performed. After data normalization, only data with a signal-to-noise ratio ≥3 were used for further analysis using the NanoString GeoMx DSP Interface provided online. This interface was used to visualize the region of interest in the epifluorescent images, perform principal component analysis, hierarchical clustering, and generate heatmaps and volcano plots with statistical analysis. Differential protein expression with p-value < 0.05 was considered significant.

### Immunohistochemistry

Zinc-formalin-fixed tissues were stored in 70% ethanol prior to paraffin embedding and 5 µm sectioning. The sections were deparaffinized using HistoChoice Clearing Agent (#H103, VWR, USA) and rehydrated using a gradient of decreasing ethanol concentrations, then in water. Some of the sections were treated with 0.45% H2O2 for 15 min before antigen retrieval by boiling in a citrate-based (pH 6.0) or Tris-based (pH 9.0) antigen unmasking solution from Vector Laboratories, USA. After antigen retrieval, the sections were washed with DPBS containing 0.1% Tween 20 (DPBST) and then blocked with Dulbecco’s Phosphate-Buffered Saline containing 10% normal donkey serum, 0.01% Tween 20, 0.3 M Glycine for 1 hr at room temperature. Next, sections were incubated at 4 °C overnight with the primary antibody diluted in the blocking buffer. Avidin/Biotin blocking kit (#SP-2001, Vector Laboratories) was used to block biotin, biotin receptors, and avidin-binding proteins on tissue sections. After washing with DPBST, the sections were incubated for 1 hr with diluted (1:500) biotin-conjugated secondary antibody in blocking buffer. The sections were washed again with DPBST and incubated with diluted peroxidase-conjugated avidin-biotin complex (PK-6100, Vector Laboratories) for 30 min. After washing with DPBST, the sections were stained with ImmPACT DAB Substrate, Peroxidase (#SK-4105, Vector Laboratories), which stained the identified antigens brown. For staining a second antigen in the same section, the same procedure was followed for blocking, primary, and secondary antibody treatment, but streptavidin-conjugated Alkaline Phosphatase (#SA-5100-1, Vector Laboratories), and the antigens were identified by blue coloration using Vector Blue Alkaline Phosphatase (Blue AP) Substrate Kit (#SK-5300, Vector Laboratories). Some ImmPACT DAB-stained sections were further stained for nuclei with hematoxylin, followed by treatment with a bluing solution (70% ethanol containing 0.45% ammonium hydroxide). DAB-hematoxylin-stained sections were dehydrated before mounting on a permanent mounting medium (#H5600, Vector Laboratories). ImmPACT DAB and Blue AP double-stained sections were mounted with an Aqueous Mounting Medium (#BUF058B, BIO-RAD). Masson’s trichrome staining was performed using the standard method. Images were acquired using a Laxco LMI 6000 Series Inverted Microscope. For whole-slide images, images were acquired at 4x magnification and were combined using the ‘stitch’ function Laxco SeBaView software. Using Masson’s trichrome-stained slides, two-dimensional areas of adipocytes were determined using ImageJ software. Some of the whole-slide images were acquired at 20x using MoticEasyScan.

For immunofluorescence staining, the H2O2 treatment step was excluded, and fluorescent-conjugated secondary antibodies were used. Prior to adding a glass coverslip, ProLong Gold Antifade Mounting Medium with DNA Stain DAPI (#P36931, ThermoFisher Scientific) was added to the sections. Fluorescent images were acquired using an LSM900 confocal microscope, and images were prepared using Zeiss ZEN 3.8 software.

### Autoantibody quantification by an antigen array

For quantification of autoantibodies in the frozen specimens, we collected 20-30 mg pieces of tissue at multiple sites of the specimen, for a total of 200 mg. Next, the tissues were homogenized in 500 µl RIPA buffer with a protease inhibitor and centrifuged to collect clear supernatant. Next, the lysates were diluted with RIPA buffer plus protease inhibitor to a total protein concentration of 945 µg/ml. The lysates were shipped to GeneCopoeia, USA, to perform OmicsArray on the Systemic Autoimmune-associated Antigen Array (PA001-V2), a 120-plex protein microarray designed to detect IgG autoantibodies in systemic autoimmune diseases such as lupus, Sjögren’s, and rheumatic diseases.

To determine autoantibodies produced by the AAA tissues, DPBS-washed freshly collected AAA tissue was cut into ∼ 5 mm pieces on DMEM medium under sterile conditions. One piece was added to a well in a 6-well plate containing 3 mL of Gibco Opti-MEM Reduced Serum Medium (ThermoFisher Scientific), then cut into ∼1 mm pieces and cultured for the next two days. As a control, only the culture medium was added to an empty well. After culture, we collected the supernatants, centrifuged them to remove debris, and shipped them to GeneCopoeia, USA, for OmicsArray PA001-V2.

Initial data analysis was performed by GeneCopoeia. Before the assay, tissue lysates were pretreated with DNase I to remove free DNA, then diluted to 1:10 in PBST buffer. Each array also contains 4 serial dilutions of Ig control and 4 serial dilutions of anti-Ig control. The net signal intensity of each antigen in each sample was normalized to the mean signal intensity of the Ig control to obtain the normalized signal intensity for each antigen, which was used to compare autoantibody levels.

### Statistical Analysis

The data were analyzed using GraphPad Prism 8 and Excel and presented as means + SEM. Significant differences in individual autoantibody levels between AAA and control aortas were assessed using multiple unpaired t-tests with Welch correction. Differences between the groups were considered significant when the p-value was < 0.05.

## RESULTS

### Identification of vascular adipocytes using snRNA-seq

To characterize the cellular makeup of human AAA, we performed snRNA-seq using 10x Genomics on frozen AAA specimens from four patients (3 males and one female). After removing doublets and regressing out mitochondrial genes, we selected 2773 nuclei for downstream analysis.

Data visualization and cellular clustering identified 15 clusters (Figure 1A and Table S1). Cluster phenotypes were assigned using previously published highly expressed marker genes^14^ and KEGG pathway analysis of significantly expressed genes (adjusted p-value <0.05) in each cluster (Figure S1). Figure 1B presents the expression of the top three identifying genes as a heat plot. The identifiable clusters are: two smooth muscle cells (SMCs) (21.2%), one endothelial cell (EC) (15.4%), one epithelial/endothelial (5.6%), four fibroblasts (32.7%), monocyte/macrophages (4.5%), B cells (4.6%), plasma cells (1.3%), NK/T cells (4.3%), neuronal cells (4.3%), endothelial cells (EC3, 4%), and adipocytes (2.1%). These percentages of cell populations align with a previously published report on scRNA-seq of human AAA^14^. Interestingly, based on the gene expression profile, both SMC populations are closely related and represent contractile phenotypes (Figure S1). They are identified by key genes *ITGA8*, *MYH10*, *LTBP1*, and *MYH11* for SMC1, and *DES*, *RGS6*, *TBX2*, and *MYH11* for SMC2 (Figure 1B).

**Figure 1.**
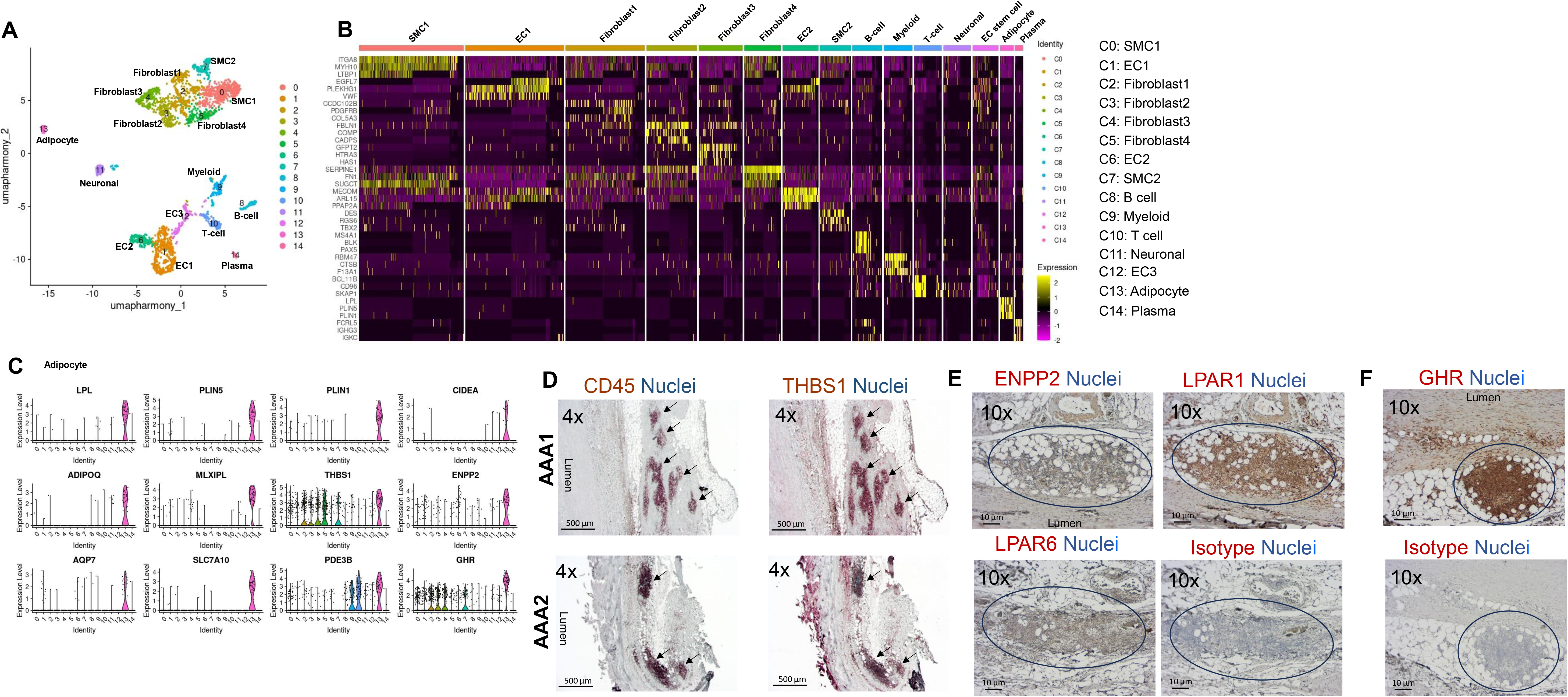
Identification and characterization of vascular adipocytes in human AAAs. **A**, Uniform Manifold Approximation and Projection (UMAP) plot showing 14 cellular clusters from snRNA-seq of aneurysm specimens from 4 patients. **B**, Heat plot illustrating the top three marker genes for each cluster, except clusters 11 and 12. **C**, Violin plots showing comparative expression of genes highly expressed in adipocytes. **D**, Immunohistochemical staining of serial cross-sections showing co-localization of CD45+ cellular clusters with thrombospondin 1 (Thbs1) in AAA from 2 patients out of 6, scale: 500 µm. **E**, Representative (n=6 patients) immunohistochemical staining of serial cross-sections showing abundance of ENPP2, LPAR1, and LPAR6 in AAA from 1 patient, scale: 50 µm. **F**, Representative (n=6 patients) immunohistochemical staining of serial cross-sections showing abundance of GHR in AAA from 1 patient, scale: 50 µm. In both E and F, background staining was identified by using an isotype control primary antibody.

We excluded clusters 11 and 12 from the heat plots. This is because cluster 11 represents neuronal cells, which have low expression levels of identifying genes^15,16^, and hence is presented as a violin plot in Figure S2. Cluster 12 has multiple mitochondrial genes, possibly because mitochondria remained fused to the nuclei, forming nucleus-associated mitochondria (NAM)^17^. Typically, NAM forms because of acute cellular mechanical confinement, and such structures promote cellular plasticity^18^. Because this cluster is close to endothelial cells, we assigned it to endothelial cells (EC3).

Cluster 13 represented adipocytes with the top 3 selected markers *LPL*, *PLIN5*, and *PLIN1* (Figure 1B). Pathway analysis revealed genes enriched for ‘Metabolic pathways’, ‘PPAR signaling pathway’, and ‘Regulation of lipolysis in adipocytes’ (Figure S1). We rigorously validated cluster 13 as adipocytes, as vascular SMCs may differentiate into adipocyte-like SMCs by overexpressing *EBF2* and *FABP4*^19^. In addition, vascular macrophages accumulate lipids, forming foam cells^20^. In particular, macrophages in human AAAs express common adipogenic marker proteins such as C/EBPβ, KLF-5, and PPAR-γ^21^. Therefore, we used additional adipocyte-specific marker genes such as *CIDEA*, *ADIPOQ*, and *MLXIPL* (Figure 1C) to confirm that cluster 13 represents adipocytes.

### Bioinformatics analysis of the roles of vascular adipocytes

Adipocytes are primarily considered a storehouse of energy in the form of fats. In obesity, some adipocytes in visceral white adipose tissue become proinflammatory and attract lymphoid and myeloid cells^22^. Multiple roles of PVAT have been described for the mouse AAAs^23^. However, it is not clear whether adipocytes in human AAA play a role in inflammation or simply act as bystanders. Using bioinformatic approaches on our snRNA-seq data, we inferred the possible roles of these vascular adipocytes based on their gene expression patterns and confirmed the spatial presence of the proteins by immunohistochemistry.

#### Transcriptional programs in the vascular adipocytes

Gene Ontology and Kyoto Encyclopedia of Genes and Genomes (KEGG) pathway analyses of significantly expressed genes (p-value <0.05) showed significant enrichment for genes involved in lipid droplet formation, fatty acid metabolism, regulation of lipolysis in adipocytes, lipid storage, the PPAR signaling pathway, the AMPK signaling pathway, and the Insulin signaling pathway (Figure S1). Other selectively and highly expressed genes in these adipocytes are: *THBS1, ENPP2, AQP7, SLC7A10, PDE3B*, and *GHR* (Figure 1C). We obtained functional annotations for these genes in humans from proteinatlas.org and provide them in Table S2.

Importantly, vascular adipocytes were the primary source of *thrombospondin 1* (*TSBS1*) (Figure 1C), a matricellular protein supporting both haptotaxis and chemotaxis potential^24,25^. In fact, THBS1 protein was highly concentrated in leukocyte (CD45+) clusters (Figure 1D). Additionally, these adipocytes were the primary source of *ENPP2*, also known as autotaxin, a secreted lysophospholipase D, that hydrolyses lysophospholipids in the extracellular matrix to lysophosphatidic acid, a bioactive lipid molecule that promotes cell proliferation, survival, migration, angiogenesis, inflammation, and fibrosis^26^. In our snRNA-seq data, SMC1 and myeloid cells highly express the lysophosphatidic acid-sensing receptors *LPAR1* and *LPAR6*, respectively. Immunohistology confirmed colocalization of ENPP2 with vascular adipocytes and TLSs (Figure 1E). Next, we tracked the protein-level expression of two highly expressed lysophosphatidic acid-sensing receptors, *LPAR1* and *LPAR6,* and found both were highly expressed in TLS microenvironment (Figure 1E). Another highly expressed gene, *SLC7A10,* regulates the fate of white and beige/brown adipocytes^27^.

Analysis of expressed transcriptional programs using geneontology.org revealed that these vascular adipocytes are distinct in having high adipocyte differentiation, *de novo* lipogenesis, lipid droplet expansion and remodeling, high lipid turnover (uptake and oxidation), high insulin sensitivity, and extracellular remodeling ability (Table 1). Thus, vascular adipocytes, by producing haptotactic and chemotactic factors, can attract immune cells and provide a niche for immature TLS formation in the AAAs.

**Table 1:**
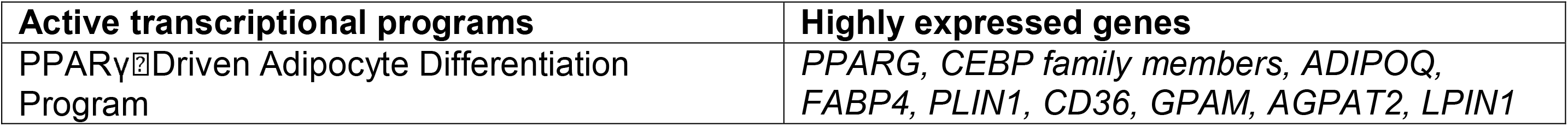
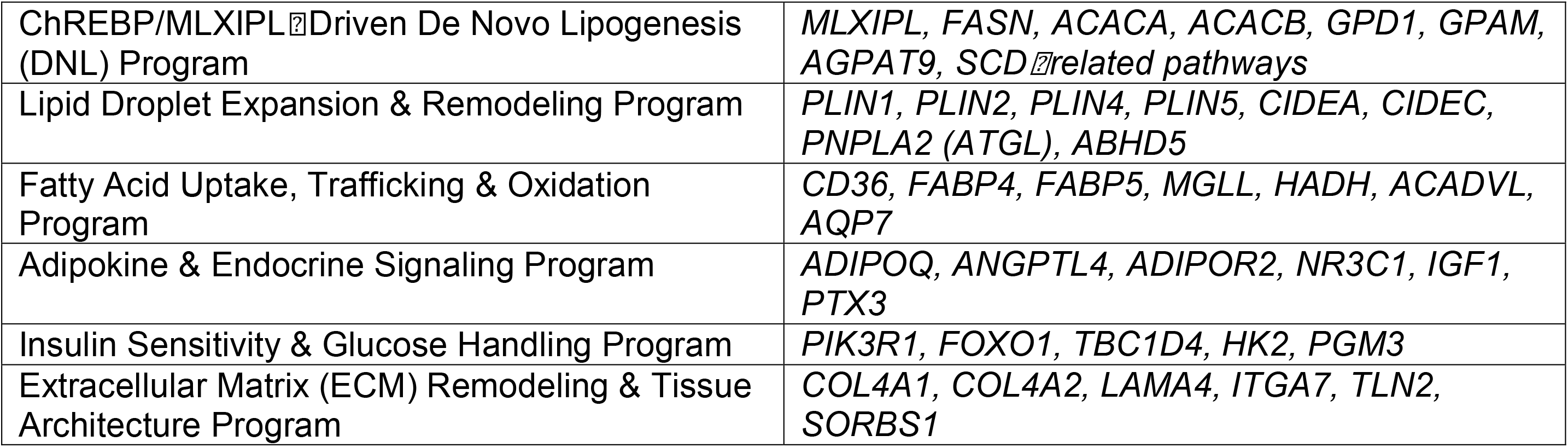

An upstream element that activates the ‘PPARγ Driven Adipocyte Differentiation Program’ is *STAT5A*^28^, which is selectively highly expressed by vascular adipocytes. A critical role of STAT5A has already been identified in a murine study in which STAT5A overexpression differentiated non-precursor cells into adipocytes, and fibroblasts overexpressing STAT5A, when injected into mice, differentiated into functional adipose tissue^28^. Interestingly, STAT5A is activated by the growth hormone receptor (GHR), whose transcript is also highly selectively expressed in vascular adipocytes (Figure 1C). Immunohistochemistry confirmed that vascular adipocytes express the protein GHR in the adipocyte-TLS microenvironment (Figure 1F). This suggests that vascular adipocytes are a potential target for Growth Hormone (somatotropin), which is secreted by the pituitary gland.

#### Vascular adipocyte and leukocyte interactions

To further discover what molecules vascular adipocytes express to establish interactions with other cells, we used the cell-cell interaction packages scDiffCom and CellChat^29^ on our snRNA-seq data. As expected, scDiffCom (600 interactions) (Figure S3, Table S3) mapped more cellular interactions compared to CellChat (105 interactions) (Figure 2A, Table S3). We focused on CellChat as it provided more comparative information. Circle plots show the number of active interactions and communication strength, highest for SMC1 (C0) (Figure 2A), a cluster with a high number of cells (filled circles at the nodes). A plot of incoming and outgoing interaction strength in Figure 2B shows that Fibroblast4 (C5) is at the top right, suggesting this cluster is highly influential, with high signal sending and receiving.

**Figure 2.**
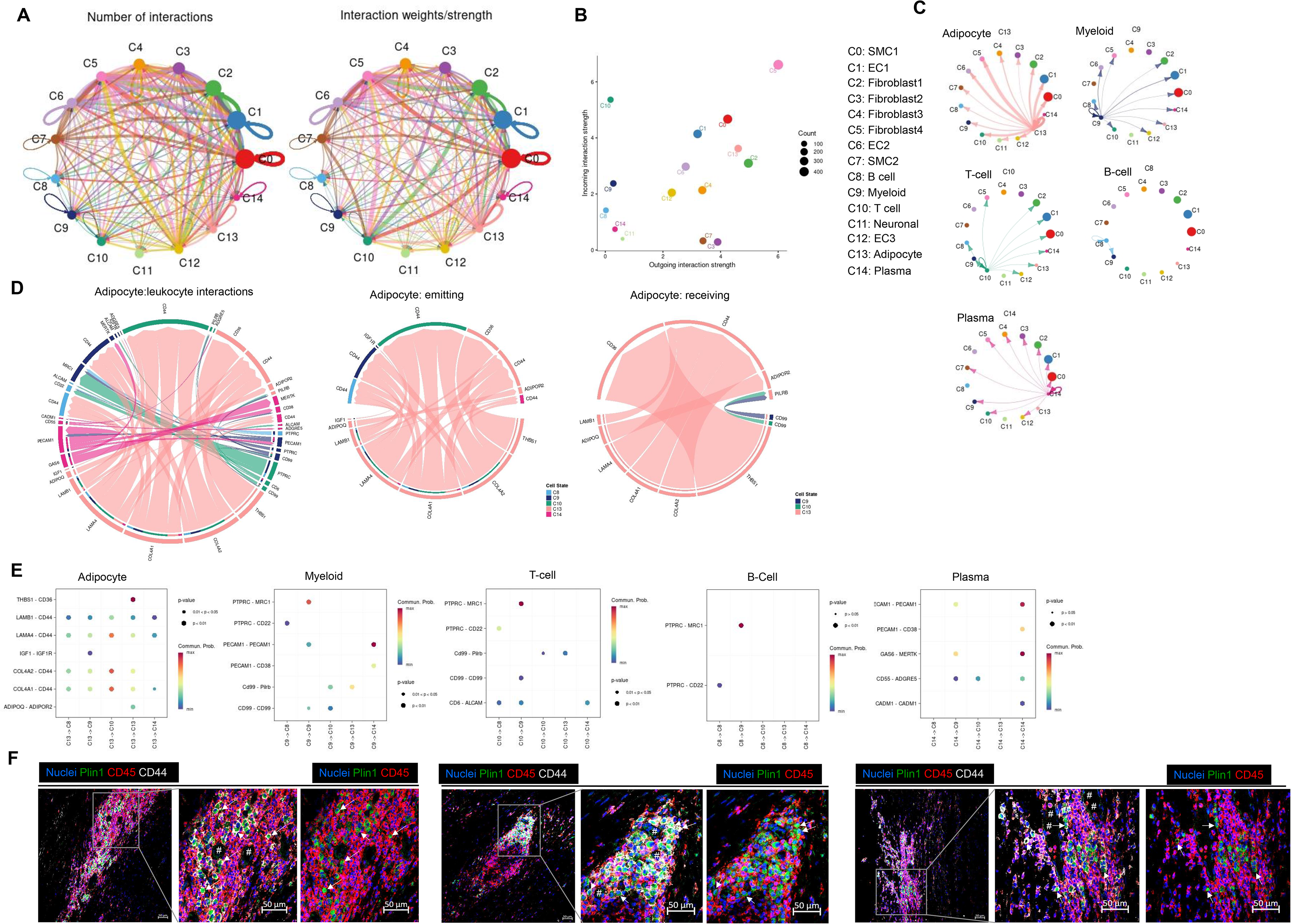
CellChat analysis of the probability of cell-cell interaction among cellular groups identified by snRNA-seq, with immunohistochemical confirmation of CD44 expression. A, Circle plot showing the probability of the number of active interactions (left) and communication strength (right) among the 14 cell groups identified in the human snRNA-seq of human AAAs. Nodes represent cell groups, and circle diameter represents the number of cells within the group. The line thickness is proportional to the total number of interactions in ‘left’ and the interaction strength in ‘right’. **B**, A scatter plot of incoming and outgoing interaction strength among the 14 cell groups. Dot color indicates cell type, and dot size indicates the number of cells. Top right: highest communication, with high signal sending and receiving, and bottom left: lowest communication. **C**, Outgoing signals from selected cell groups to the other 14 cell groups. **D**, Chord diagram showing ligand-receptor pairs and their weight for total adipocyte-immune cell interaction, followed by adipocyte emitting and receiving signals. **E**, Comparison of the significant ligand-receptor pairs between one cell type to other selected cell types. Dot color represents communication probability, and dot size denotes the p-value. **F**, Confocal images of human AAA cross-sections showing nuclei (DAPI, blue), perilipin 1 (Plin1, green), hyaluronan receptor (CD44, red), and leukocytes (CD45, white). ‘#’ indicates adipocytes identified by empty circular/oval spaces, and ‘arrow’ indicates Plin1+ CD44+ but CD45-cells in leukocyte aggregates (n=6 patients). Scale: 50 µm.

Neuronal (C11) and plasma cells (C14) are placed at the bottom left, indicating low communication. Interestingly, adipocytes (C13) were placed near the top right, suggesting they actively communicate with the localized microenvironment within the AAA. Furthermore, adipocytes highly express genes whose protein products can establish communication with the remaining 13 cellular clusters (Figure 2C) and with other adipocytes.

As per CellChat analysis, among the leukocytes, the myeloid cells did not send any signals to C3, C4, C7, and C11; the T-cell/NK cell populations did not send any signals to C3, C4, C6, C7, and C11; and the plasma cells did not send any signals to C6, C8, and C11 (Figure 2C). B cells sent signals only to myeloid cells (C9), plasma cells to all cell types except neuronal cells, and T cells to B cells, plasma cells, myeloid cells, adipocytes, SMC1, EC1, EC3, fibroblast1, and fibroblast4. To identify potential interactions between vascular adipocytes and immune cells, we analyzed ligand-receptor pairs between adipocytes and B cells, T cells/NK cells, plasma cells, and myeloid cells. The signals emitted from adipocytes to other immune cells were *LAMB1 - CD44*, *LAMA4 - CD44*, *COL4A2 - CD44*, and *COL4A1 - CD44* (Figure 2D and E). According to genecards.org, *LAMB1, LAMA4, COL4A2*, and *COL4A1* encode structural basement membrane proteins. *CD44* encodes the cell-surface protein CD44, also known as a hyaluronan receptor. CD44 expression on inflammatory leukocytes promotes cellular adhesion^30,31^. We confirmed CD44 protein expression on leukocytes (CD45+) using immunohistochemistry (Figure 2F). Unexpectedly, we found Plin1+ CD44+ but CD45-cells in leukocyte aggregates (Figure 2F), suggesting these cells are adipocyte progenitor cells or dedifferentiated adipocytes^32^.

Together, these results suggest vascular adipocytes as a potential site for inflammatory leukocyte recruitment within AAA.

### TLSs were associated with vascular adipocytes in human AAAs

To determine the spatial organization of vascular adipocytes and TLSs in AAA tissues, we performed immunohistological analyses of AAAs and control abdominal aortas from organ donors. Our cohort included AAA specimens from 13 patients undergoing elective repair (2 with occlusive AAA), 1 popliteal artery aneurysm (PAA), and control abdominal aortas from 4 organ donors (Table S4). We obtained AAA tissue sections from one AAA patient from OriGene, MD, USA. Altogether, we had cross-sections from 14 AAAs, 1 PAA, and 4 control abdominal aortas. Masson’s trichrome staining of the aortic cross-sections demonstrated collagen deposition (blue coloration) in the intima, media, and/or adventitia of the aneurysm tissues (Figure 3A and Figure S4). The non-aneurysmal controls showed varying levels of collagen: control1 had high collagen in the intima-media, control2 had partial collagen in the adventitia, and control3 and control4 had collagen both in the intima and adventitia (Figure 1A and Figure S5). Thrombus at the luminal side was prominent in AAA5, AAA10, AAA11, control3, and control4. Atherosclerosis, identified by cholesterol crystals (elongated/rhomboid, sharp-edged empty spaces), was prominent in AAA4. Furthermore, alpha-smooth muscle actin (aSMA) staining showed significant destruction of medial smooth muscle cells compared with non-aneurysmal controls (Figure 3B and Figure S6A and B). We did not detect the endothelial layer (CD31+) in the intima of either the AAAs or controls. However, we found strong aSMA and CD31 staining in the vasa vasorum of both AAAs and controls. Interestingly, localized aSMA and CD31 staining around the advanced TLSs, and CD31 staining within the TLS (Figure 3B and Figure S6A), suggest close interactions among endothelial cells, smooth muscle cells, and TLSs. These results established our aneurysmal and control specimens for further spatial analysis of vascular adipocytes and TLSs.

**Figure 3.**
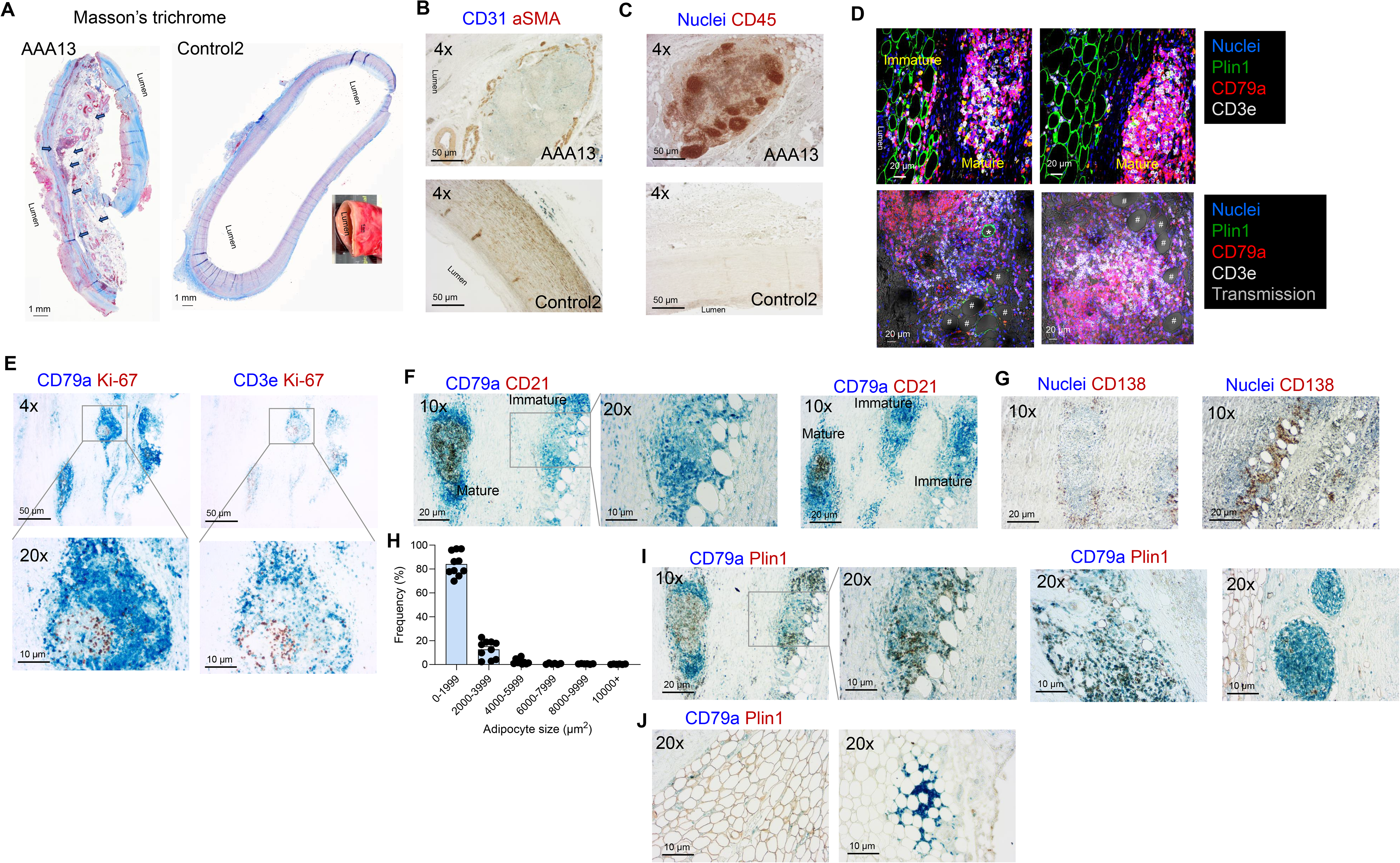
Histological analysis of vascular adipocytes and TLSs in human AAAs. **A**, Whole-slide images of Masson’s trichrome staining of the cross-sections of a AAA and a control abdominal aorta showing unilocular adipocytes (arrows) (n=15 patients and 4 control abdominal aortas). The inset in the control abdominal aorta shows unmanipulated tissue used for paraffin embedding, followed by cross-sectioning. Blue: collagen; red: muscle, cytoplasm; and black: nuclei. **B**, Immunohistochemical staining of cross-sections of a AAA and a control abdominal aorta showing endothelial cells (CD31, blue) and smooth muscle cells (α-smooth muscle actin, aSMA, brown) (n=6 patients and 4 control abdominal aortas). **C**, Immunohistochemical staining of cross-sections of a AAA and a control abdominal aorta showing nuclei (Hematoxylin, blue) and clusters of immune cells (CD45, brown) (n=15 patients and 4 control abdominal aortas). **D**, Multiple confocal images of AAA cross-sections showing adipocytes (Plin1, green) and immature and mature TLSs with nuclei (DAPI, blue), B cells (CD79a, red), and T cells (CD3e, white). Transmission images were overlaid in the bottom two images to show Plin1+ adipocytes (#) and Plin1-adipocytes (*) in the TLSs (n=6 patients). **E**, Co-staining of serial AAA cross-sections for B cells (CD79a, blue)-Ki-67 (brown), and T cells (CD3e)-Ki-67 (brown) to show dividing lymphocytes in the TLSs (n=7 patients). **F**, 10x and 20x images of AAA cross-sections showing B cell (CD79a, blue) clusters with dendritic cells (CD21, brown) representing mature TLS, and B-cells without dendritic cells representing immature TLS (n=11 patients). **G**, AAA cross-sections stained with nuclei (hematoxylin) and plasma cells (CD138, brown) showing plasma cells around the border of a TLS (left panel) or scattered around vascular adipocytes (right panel) (n=10 patients). **H**, Quantification and size distribution of adipocyte areas (2-dimensional) determined using ImageJ, using 3230 adipocytes in Masson’s trichrome-stained AAA cross-sections from 9 patients. Values were expressed as means. **I**, B cell (CD79a, blue) clusters and adipocytes (Plin1, brown) staining of AAA cross-sections to show media-adventitia localized adipocyte clusters (n=6 patients). **J**, B cell (CD79a, blue) clusters and adipocytes (Plin1, brown) staining of AAA cross-sections to show a continuous layer of adipocytes in the periadventitial region (n=6 patients). Scale bars are shown in each image.

We initially identified TLSs by leukocyte clusters (CD45) (Figure 3C), then stained for B cells (CD79a) and T cells (CD3e) (Figure 3D). We broadly classified TLSs into three categories: 1) non-TLS aggregates, identified by scattered aggregates of B cells or T cells, 2) non-classical/immature TLS, identified by randomly organized clusters of B cells and T cells, and 3) classical/mature TLS, identified by organized clusters of B cells and T cells with at least one germinal center, CD21+ dendritic cells, or actively dividing (Ki-67+) lymphocytes, as described before^33^ (Figure 3D-F). Plasma cells are localized at the border of the mature TLSs or scattered near the vascular adipocytes (Figure 3G). Of the 15 aneurysm specimens, 10, including PAA6, had TLSs in the adventitia. Non-TLS aggregates and immature TLSs were identified in all 10 specimens. Additionally, AAA1, AAA13, AAA14, oAAA15, and AAA16 showed mature TLSs.

Vascular white adipocytes were identified as large (>20 µm), unilocular, lipid-filled cells (characterized by empty circular/oval spaces) that stained positive for perilipin 1 (protein product of *PLIN1*) (Figure 3D). Perilipin 1 is a lipid droplet-coating protein and is required for maintaining futile lipolysis in healthy adipocytes. Out of 15 aneurysm specimens, 13 contained adipocytes, and adipocytes were associated with TLSs in 10 of them (Supplemental Table 2). To determine adipocyte size, we analyzed 3230 adipocytes from aneurysm cross sections from 9 patients. The adipocytes had a median 2-dimensional size of 997 ± 49 µm^2^ (Figure 3H), corresponding to a diameter of 35.6 µm. This adipocyte size range is smaller than that of subcutaneous (2679±795 µm^2^) and visceral adipocytes (2100±719µm^2^) determined by histological methods in previous studies^34,35^. Additionally, we had three interesting observations: (i) the 3 non-aneurysmal controls and 3 out of 15 AAA specimens had few to no adipocytes (Figures S4 and S5), suggesting that adipocytes are not common in human abdominal aortas. (ii) Some of the adipocytes lacked Plin1 staining (Figure 3D and I). Plin1-negative adipocytes are considered dysfunctional and proinflammatory adipocytes^36^. Additionally, some Plin1+ cells, which were confirmed by both immunofluorescence and diaminobenzidine staining, were present within the TLSs (Figure 2F, 3D and I). (iii) Immature TLSs surround some of the adipocytes in 10 out of 15 aneurysm specimens (Figure 3D), similar to a crown-like structure in white adipose tissues^37^, suggesting these adipocytes are proinflammatory.

Interestingly, unlike murine PVAT, some isolated and clustered adipocytes were localized farther from the perivascular region and in the medial-adventitial region, suggesting these adipocytes are generated in the AAA (AAA13 in Figure 3A, Figure 3D, and I). Additionally, in Figure S4, AAA2, PAA6, and oAAA12, and in Figure S5, Control1, show a heterogeneous layer of adipose tissue in the perivascular region. Controls 2, 3, and 4 in Figure S5 show a few clusters of adipocytes in the perivascular region. Control 4 has adipose tissue, but it is localized outside of the perivascular region. These results suggest that vascular adipocytes are formed at least in two different ways in abdominal aortas: (i) a layer of adipocytes in the perivascular region, which is generally known as PVAT, and (ii) isolated adipocytes or clusters of adipocytes in the medial-adventitial region.

Altogether, these results confirm the previously reported positive association of AAA with TLS^38^ and AAA with adipocytes^39,40^. Here, we discovered that TLSs were associated with vascular adipocytes. Additionally, adipocytes are not a normal constituent of non-aneurysmal human abdominal aortas and form in human AAA tissues.

### High-resolution spatial transcriptomics analysis of TLSs in human AAAs

The snRNA-seq data lacks critical pathological spatial information such as the cellular organization of SMCs, endothelial cells, T cells, and B cells in TLS. Therefore, we performed high-definition spatial transcriptomic profiling using Visium HD, which generated gene expression data from 2 µm x 2 µm barcoded squares. Data from 8 µm (4 x 4) bins and 16 µm (8 x 8) bins showed that 16 µm outperformed 8 µm, with reduced data sparsity, higher mean UMI counts/bin, and better representation of snRNA-seq clusters (Figure 4A and Figure S7).

**Figure 4.**
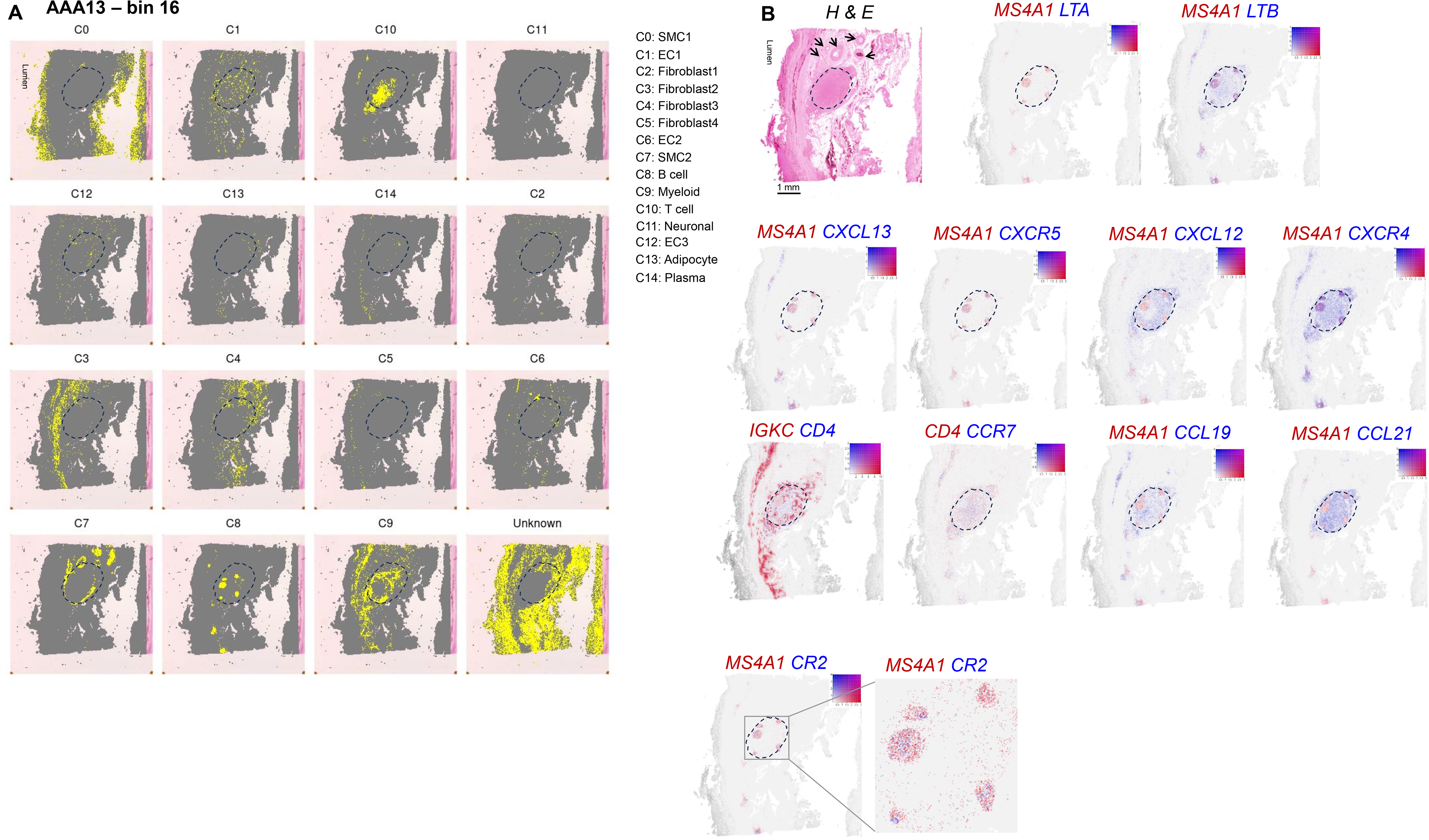
Visium HD spatial transcriptomics on human AAA13. **A**, 16 µm (8 x 8) bins of Visium HD spatial transcriptomics on AAA13 showing localization of all 14 cellular clusters obtained from snRNA-seq on the AAA cross-section. **B**, Hematoxylin and Eosin (H&E)-stained image of AAA13 used for Visium HD spatial transcriptomics is shown, followed by spatial images showing gene expression patterns to show co-localization of B cells (*CD79A1*) with various chemokines and chemokine receptors, followed by plasma cells (*IGKC*), CD4+ T cells (*CD4*), and CD21 (*CR2*). The images were generated using the Loupe browser, and gene expression levels are shown. The scale bar in the H&E image is 1 mm, and arrows indicate vasa vasorum.

**Table 2:**

| Bin Metrics Overview |  |  |  |
| --- | --- | --- | --- |
| Specimen id | Bin Size (µm) | 8 µm | 16 µm |
| A13 | Number of Bins Under Tissue | 375,997 | 95,749 |
|  | Mean UMI Counts per Bin | 66.3 | 261.1 |
|  | Mean Genes per Bin | 54.8 | 193.8 |
| A1 | Number of Bins Under Tissue | 159,157 | 40,730 |
|  | Mean UMI Counts per Bin | 17.9 | 70.4 |
|  | Mean Genes per Bin | 13.3 | 48.1 |

Projection of our snRNA-seq data onto the spatial data revealed that the SMC1 and SMC2 populations, which were phenotypically indistinguishable from each other in snRNA-seq data, were localized in the spatial data: SMC1 (C0) to the intima-medial region and SMC2 (C7) specifically to the small vessels within the aorta (vasa vasorum) for specimen A13 (Figure 4A). Additionally, SMC2 partly encircles TLSs, similar to a lymph node capsule. Specimen A1 also positioned both SMC populations as spatially distinct; in particular, SMC2 was colocalized with an advanced TLS and vasa vasorum (Figure S7B). The spatial maps also placed ECs, fibroblasts, adipocytes, and immune cell populations distinctly in both specimens (Figure 4A).

Both EC1 (C1) and EC2 (C6) cells were localized throughout the AAA section. In particular, the EC1 cluster, enriched for genes that promote leukocyte recruitment and organization, such as *ICAM1*, *SELE*, *CD34*, and *PECAM1*, is localized both within and outside the TLSs. In our snRNA-seq data, the fibroblast 4 cluster (C5) highly expressed *VCAM1*, and the myeloid cluster (C9) highly expressed *ICAM1* (Table S1). These adhesion molecules are critical for leukocyte attachment and immobilization at the site of aortic inflammation.

After leukocyte immobilization, various cytokines and chemokines direct the spatial organization of B cells and T cells to form immature TLS, which mature into classical TLS. During TLS formation, the lymphotoxin α (*LTA*) and lymphotoxin β (*LTB*) heterotrimer synthesized on the surface of TLS inducer (LTSi) cells binds to LTβR expressed on TLS organizer (TLSo) and promotes cell surface expression of VCAM1 and ICAM1. The interaction between TLSi and LTSo leads to production of the homeostatic chemokines CXCL13, CCL19, and CCL21^41,42^. We localized the selective expression of *LTA* and *LTB*, as well as *CXCL13*, *CCL19*, and *CCL21,* within the TLSs (Figure 4B). Interestingly, *CXCL13* and its cognate receptor, *CXCR5*, were confined to the CD20 (*MS4A1)*+ B-cell areas of TLSs, suggesting that these areas are B-cell follicles. In both AAA sections A13 and A1, multiple such B-cell follicles were clustered to form a nodular structure (dotted areas).

In secondary lymphoid organs (SLOs), primary B-cell follicles contain naïve B-cells that, after antigen and T-cell activation, transform into secondary B-cell follicles with distinct light and dark zones. CXCL13 binds CXCR5 on B cells and promotes B-cell aggregation in the light zone for efficient antigen selection, whereas CXCL12 binds CXCR4 on B cells to promote B-cell migration to the dark zone, a suitable microenvironment for immunoglobulin somatic hypermutation^41,42^. In our Visium data, while *CXCL13* and *CXCR5* expression were confined to *MS4A1+* B-cell follicles, representing centrocytes and the light zone. Additionally, *CXCR4* was highly expressed in parts of the B-cell follicle, representing centroblasts, the proliferating B-cells in the dark zone. Low *CXCR4* expression was also localized to the parafollicular areas within the nodular structure. Our snRNA-seq experiment revealed that T-cells and plasma cells primarily express *CXCR4*. Visium data revealed that strong *CXCL12* expression was confined to the parafollicular area, possibly originating from endothelial cells (Figure 3B, top image).

Additionally, the parafollicular area within the nodular structure is filled with plasma cells (*IGKC*) and *CD4+* T cells (Figure 4B). *CCL19* and *CCL21* expression was localized to the parafollicular area. In SLOs, CCL19 and CCL21 bind to CCR7 on T cells and promote organization of the T-cell zone. In agreement with this, *CCR7*-expressing T cells were localized in the parafollicular areas (Figure 4B). Additionally, *CR2* (CD21), expressed on the surface of follicular dendritic cells in B-cell follicles, efficiently traps C3d-coated antigens, increasing local antigen concentration and presenting them to antigen-specific B cells. In fact, *CR2* gene- and protein-expressing cells were found in the B-cell follicles (Figures 4B and 3F).

Altogether, these results suggest that TLSs within AAAs recapitulate SLO ontogeny.

### Human AAA TLSs are potential sites for generation of B-cell immunoreactivity

To determine B-cell subtypes present in human AAAs, we analyzed available scRNA-seq data on human AAA (GSE166676^14^). This dataset has ∼8000 cells from AAA tissues from 3 patients and control aortic tissues from 2 individuals. Data visualization and cellular clustering identified 26 clusters (Figure S8A). Among the 26 cellular clusters, the closely placed clusters 1 and 5 were B-cells based on high *MS4A1* (CD20) expression (Figure S8B). These B-cells are further identified into 5 subsets (Figure S8C, and Table S5): cluster 0: follicular B-cells (high expression level of *TCL1A*, *FCER2*, *CD79A/B*, *HLA-DR* Family, *IL4R,* and *KLF2*); cluster 1: transitional B-cells (high expression level of ribosomal proteins, long non-coding RNAs *MALAT1* and *NEAT1*, and stress-response genes *TP53INP1* and *TXNIP*); cluster 2: CD27+ memory B-cell (high expression level of *CD27*, *TNFRSF13B*/*TACI*, and mitochondrial genes); cluster 3: CD1D+ Memory B cells (high expression levels of *CD1D* with genes supporting memory phenotype *FN1*, *COTL1*, *SLC2A3*, *ACAP1*, *NCK2*, *CAPN2*, and *VPS37B*); and cluster 4: proliferative germinal center, particularly, Dark Zone (DZ) B-cells or centroblasts (high expression levels of *MKI67*, *AICDA* and *BCL6*, DNA replication & histones). Interestingly, cluster 2 highly expressed *BHLHE41*^43^, a murine B-1 B-cell marker. Cluster 15 highly expressed *MZB1, CD38, IGJ*, and BAFF receptor *TNFRSF17 (BCMA),* representing plasma blasts and plasma cells. Altogether, the presence of these B-cell subtypes suggests that germinal center reactions may proceed to plasma cell and memory cell formation within AAA tissues.

Our snRNA-seq data revealed that plasma cells highly express the immunoglobulin (Ig) heavy chain genes *IGHG1*, *IGHG3*, *IGHG4*, *IGHA1*, *IGHGP*, and *IGHM*, and the light chain genes *IGKC* and *IGLC2* (Table S1). These plasma cells also highly express plasma cell surface markers *SDC1 (CD138)* and *CD38*, and canonical transcriptional programs *PRDM1 (BLIMP 1)* – master regulator of plasma cell fate, *XBP1* – unfolded protein response, *MZB1* - an endoplasmic reticulum chaperone required for Ig production, and *IRF4* – essential for plasma cell survival. Importantly, plasma cells highly expressed *IL21R*, suggesting a skew toward IgG and IgA synthesis. As seen in the scRNA-seq data, plasma cells did not have detectable levels of *TNFRSF17* (Table S1).

Pathway analysis revealed genes enriched for ‘Protein processing in endoplasmic reticulum’ and ‘Leukocyte transendothelial migration’ (Figure S1). This suggests that the plasma cells migrated from blood – this we analyzed in further detail. There are multiple possible ways plasma cells can localize in AAA tissues: (i) directly migrated from the blood, (ii) differentiated from migrated memory B-cells, (iii) generated from local TLSs, or (iv) generated in a local TLS and migrated to another site within the tissue.

While snRNA-seq and scRNA-seq provide limited information on the source of plasma cells, our spatial transcriptomics provided additional insights. In particular, we identified clusters of plasma cells expressing *IGHG1, IGHG3, IGHA1,* or *IGHM* (Figure 5A). Among the heavy chains, *IGHG1* was highly expressed, and among the light chains, *IGKC* was higher than *IGLC* – this is consistent with our snRNA-seq data (Table S1). Using immunohistochemistry, we confirmed IgD+, IgG+, IgM+, and IgA+ cells in TLSs (Figure 5B). Interestingly, both AAA and control aortas showed strong IgG deposition, whereas only the intima of control aortas showed IgM and IgA deposition (Figure 5B). Consistent with very low *IGHE* expression, we found only a few cells positive for IgE by immunohistochemistry (Figure 5A and B).

**Figure 5.**
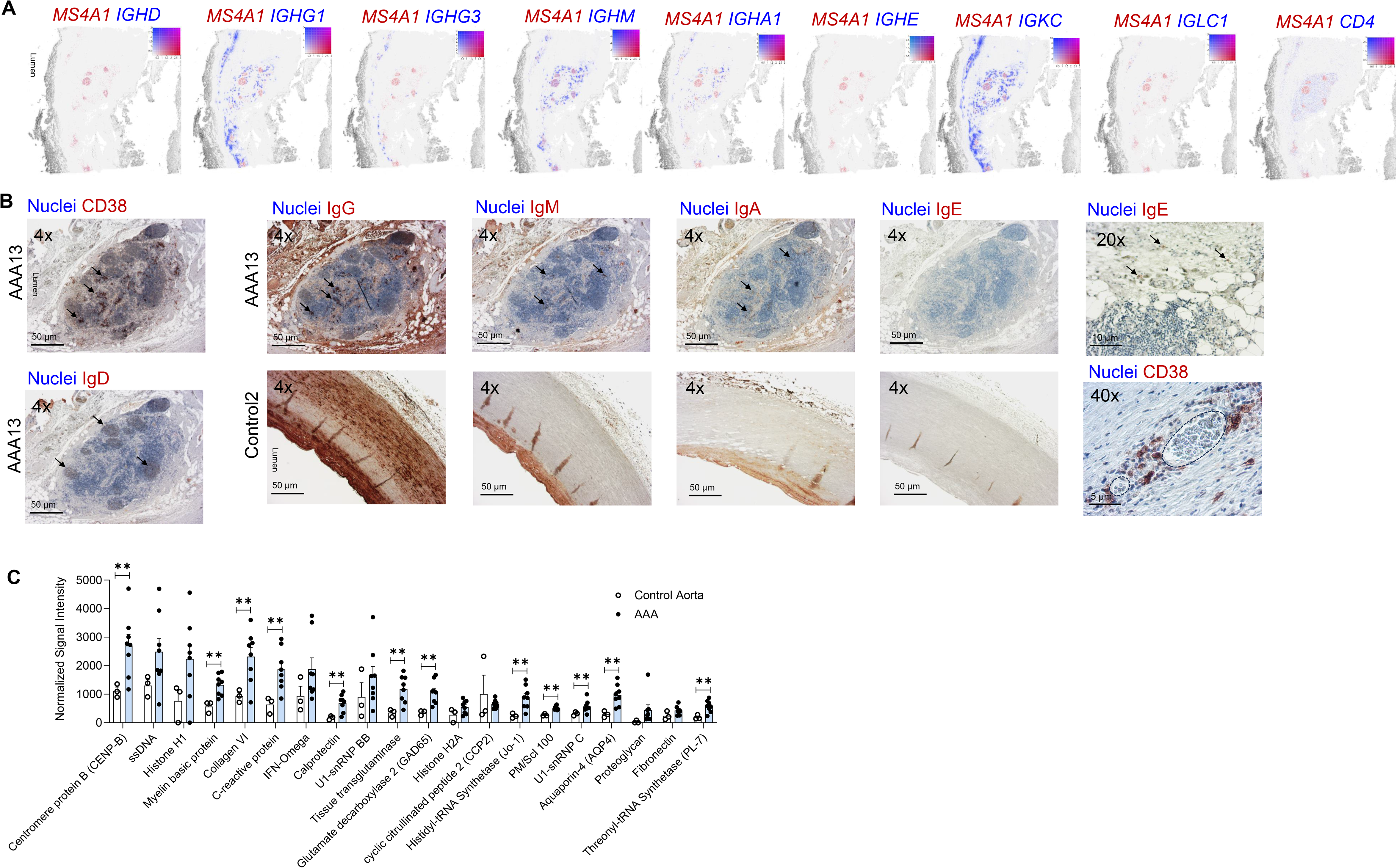
Characterization of B-cell autoreactivity in human AAAs. **A**, 16 µm bins of Visium HD spatial transcriptomics on AAA13 showing spatial images with gene expression patterns to illustrate co-localization of B cells (*CD79A1/CD20*) with various immunoglobulin heavy-chain and light-chain gene expression. CD4+ T cell (*CD4*) areas are shown. Some images repeat Figure 4 for comparative analysis of T-cell and B-cell areas. The images were generated using the Loupe browser and show gene expression levels. **B**, Serial cross-sections of AAA and control aortas were stained for CD38 (plasma cells), IgG, IgM, IgA, IgE, and IgD (brown), and for nuclei (Hematoxylin) to localize plasma cells and immunoglobulin-expressing cells (arrows) in TLSs. Representative images of AAA13 and control2 are shown. One 20x image of a non-serial section from AAA13 is presented to show cells positive for IgE (arrows). A 40x image shows clusters of plasma cells (CD38+) near a blood vessel (dotted lines) (n=6 patients for CD38, and n=14 patients and 4 control abdominal aortas for immunoglobulins). **C**, Quantification of the top 20 IgG autoantibodies in the AAA (n=8) and control abdominal aorta (n=3) lysates (protein concentration in each lysate was adjusted to 945 µg/ml). Values are expressed as means + SEM. ‘**’ indicates p< 0.01 determined by multiple unpaired t-tests with Welch correction.

In TLSs, the presence of plasma cells in close proximity to germinal centers (Figure 3G, CD138; Figure 5B, CD38) is indicative of the process in which naïve B-cells that are challenged by autoantigens enter the germinal center reaction and Ig isotype switching, and finally mature into plasma cells^33^. In this context, we localized plasma cells of various Ig isotypes near germinal centers (Figure 5A). Interestingly, we also found plasma cells in non-classical TLSs (Figure 3G), suggesting differentiation of memory B-cells into plasma cells after ligation by antigen-specific T-cells. Some plasma cells were found scattered in the intima-media and adventitial regions (Figure 5B, CD38), possibly having migrated directly from the blood or from local TLSs. Plasma cells were also localized near the vasa vasorum, suggesting their migration from the blood (Figure 5B). Thus, our results suggest that plasma cells in AAA derive from multiple sources, including local activation and differentiation of naïve B-cells in TLSs.

The presence of plasma cells with actively transcribed Ig genes in both our snRNA-seq and spatial transcriptomics data suggests that AAA tissues synthesize Ig. Because IgG genes were highly expressed, we next identified IgGs’ target antigens by performing an antigen array on tissue lysates from control abdominal aortas and AAA specimens. Interestingly, the highly enriched antigens were of both extracellular and intracellular origins (Figure 5C). Among the highly abundant top 20 autoantibodies (autoAbs), those that were significantly higher in the AAAs were against the antigens Centromere protein B (CENP-B)^44^, Myelin basic protein^45^, Collagen VI^46^, C-reactive protein^47^, Calprotectin, Tissue transglutaminase, Glutamate decarboxylase 2 (GAD65)^48^, Histidyl-tRNA Synthetase (Jo-1)^49^, PM/Scl 100^50^, U1-snRNP C^51^, Aquaporin-4 (AQP4)^52^, and Threonyl-tRNA Synthetase (PL-7)^53^, which were also found in autoimmune diseases such as lupus. Other abundant autoAbs that trended toward significance in the top 20 were ssDNA, Histone H1, IFN-Omega, U1-snRNP BB, Histone H2A, cyclic citrullinated peptide 2 (CCP2), Proteoglycan, and Fibronectin^54^.

Next, to determine whether plasma cells within the AAA tissues secrete these autoAbs, we cut a part of the freshly collected AAA tissues into small pieces, washed them to remove contaminating blood, then cut the tissues into 1 mm pieces, cultured them in a low-serum medium for 2 days, and measured Abs in the culture supernatants. Qualitative assessment of Igs using test strips identified IgG1 as the dominant antibody isotype and κ-chain as the preferred light chain (Figure S9A), aligning with our spatial transcriptomics results (Figure 4B). An antigen array of the culture supernatants revealed that the dominant IgG Ab was against Epstein-Barr nuclear antigen-1 (Figure S9B), which is identified in 95% of the US population aged ≥40^55^. AutoAbs to elastin were the most abundant in AAA tissues (reported^56^). Other autoAbs included GAD65, cardiolipin^57^, Factor B^58^, Fibronectin, Collagen VI, Type-1 angiotensin II receptor^59^, and fibrinogen S^60^, some of which were identified in AAA tissue lysates. While more studies are needed to determine whether these autoantibodies are monoreactive or polyreactive, these data suggest that human AAA lesions may be sites of local B-cell autoreactivity.

### Spatial proteomics and transcriptomics identified active ERK1/2 signaling during the maturation of aneurysmal TLS

Unlike murine models of AAA, human AAAs develop over years. Therefore, analyzing immune cell infiltration in the early stages of aortic inflammation, when non-TLS lymphocyte aggregates form, and in well-established AAA tissue, where multiple mature TLSs are present, will provide insight into critical changes in immune cells during AAA development. To this end, we performed NanoString GeoMx DSP profiling of proteins on the four non-TLS aggregates in one control aorta and four TLSs in one advanced AAA lesion (Figure 6A, both obtained from OriGene, USA) and determined cellular activation status and intracellular signaling events in macrophages (CD68+), B cells (CD20+), and T-cells (CD3+). After identifying 4 regions of interest (ROI) within each aorta, we quantified 79 targets from Immuno-Oncology Protein Panels (Table S6) within each cell type using UV-cleavable DSP tags.

**Figure 6.**
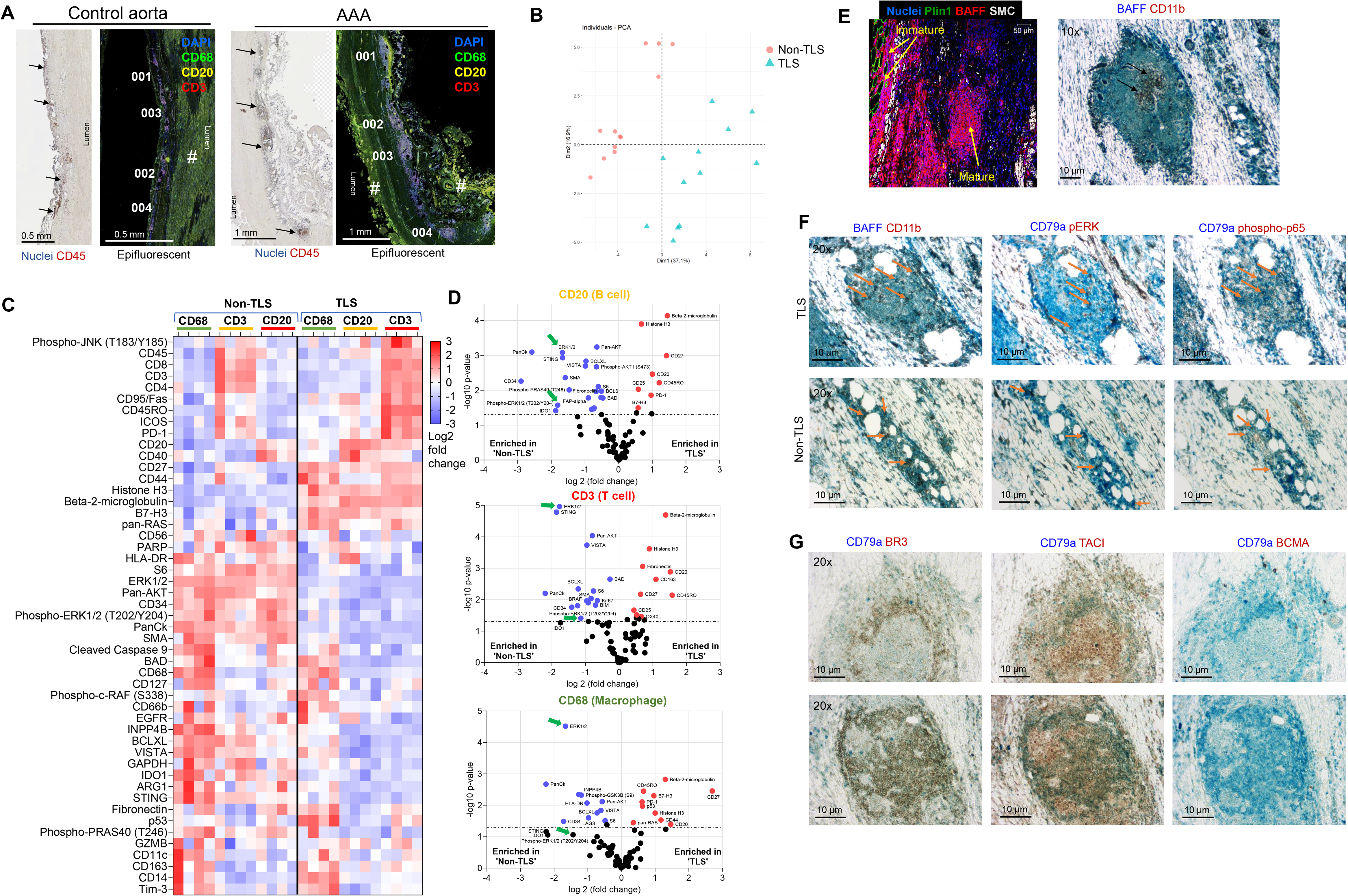
Spatial analysis of protein expression in non-TLS aggregates and TLSs. A, Whole-slide image of control aorta and AAA stained for nuclei (Hematoxylin) and immune cells (CD45), followed by epifluorescent images of serial sections showing morphology markers for B-cells (CD20, yellow), T-cells (CD3, red), and macrophages (CD68, green) used for NanoString GeoMx Digital Spatial Profiling of proteins. Based on clustering of the morphology markers, we selected 4 regions of interest (‘001’ to ‘004’) in the adventitial site of the control aorta; similarly, we selected 4 regions in the AAA slide. Each region contained B-cells, T-cells, and macrophages. Lumens are indicated. ‘#’ indicates autofluorescence. Arrows indicate the clusters of immune cells in the adventitia. **B**, Principal component analysis (PCA) of protein expression in the 4 non-TLS aggregates and in the 4 TLSs. Each symbol represents a protein expression pattern in B-cells, T-cells, or macrophages. **C**, Heat map illustrating hierarchical organization of protein expression in B-cells, T-cells, and macrophages, followed by **D**, illustrating comparative protein expression in each immune cell type in the non-TLS and TLS. The dotted line indicates the significance threshold (p <0.05). Significantly enriched proteins in non-TLS are shown as blue dots, and in TLS as red dots. Green arrows indicate ERK1/2 and phospho-ERK1/2 (T202/Y204). **E**, Left, A confocal image of an AAA cross-section showing localization of BAFF (red) in an immature and a mature TLS microenvironment. Nuclei (DAPI), adipocytes (Plin1, green), and smooth muscle cells (aSMA, white), n=6 patients. Right, A light microscopy image of an AAA cross-section showing co-localization of BAFF (blue) with myeloid cells (CD11b, brown) (n=10 patients). **F**, Serial cross-sections of AAAs showing TLS (top panels) and non-TLS aggregates (bottom panels) stained for BAFF (blue) and myeloid cells (CD11b, brown); B-cells (CD79a, blue) and phospho(Y202/Y204)-ERK1/2 (brown); and B-cells (CD79a, blue) and phospho(S536)-p65 (brown) (n=10 patients). Some double-positive staining is indicated with arrows. **G**, Serial cross-sections from two AAAs (top and bottom) showing co-localization of BAFF receptors BR3, TACI, and BCMA (brown) with B-cells (CD79a, blue). Scale bars are shown in each image; n = 7 patients.

Principal component analysis clearly separated the protein expression patterns in non-TLS from TLS (Figure 6B). Notably, the heatmap (Figure 6C) and volcano plot (Figure 6D) revealed that TLSs were enriched for B-cells expressing CD45RO and CD27, suggesting activated, highly mutated human germinal center B-cells and somatically hypermutated antigen-specific B-cells capable of producing immunoglobulins, respectively. Further, TLSs were enriched for CD45RO-expressing T-cells, suggesting memory T-cells. Interestingly, some of the T-cells express CD20, which is possibly acquired from B-cells *via* trogocytosis, and such T-cells are highly inflammatory effector cells^61^. High CD27 expression in macrophages may arise from engulfed T-cells or B-cells. In the non-TLSs, notable signals were from ERK1/2, phospho-ERK1/2 (T202/Y204), pan-Cytokeratin (PanCk), CD34, and the anti-apoptotic protein BCL-XL from macrophages, B-cells, and T-cells. Expression of PanCK and CD34 suggests that some of these cells are in their precursor stage, differentiating from hematopoietic stem cells^62–64^. Altogether, these results suggest that lymphocytes are activated for further differentiation in the non-TLSs.

ERK1/2 signaling is critical for the survival of B-cells by preventing apoptosis after antigenic stimulation *via* the B-cell receptor (BCR)^65^. BAFF induces this signaling and prevents B-cell apoptosis. Immunohistochemical staining revealed high BAFF staining in both non-TLS aggregates and mature TLSs (Figure 6E). In line with high BAFF concentration, critical BAFF and BCR signaling proteins phospho-ERK1/2 [T202/Y204] and canonical NF-kB signaling protein phospho-p65 [S536] were localized in immature and mature TLSs (Figure 6F), suggesting activation of the coupled BR3-BCR signaling pathway, which is separately mediated by BAFF and antigen cross-linking by BCR^66,67^. In agreement with this, TLSs highly expressed BR3 and TACI receptors (Figure 6G) and BCRs, that is, IgD+ cells and IgM+ cells in Figure 5B.

To identify the potential source of BAFF in AAA tissues, we analyzed BAFF transcripts in our snRNA-seq data and the available scRNA-seq data (GSE166676). Although BAFF transcript levels were low in our snRNA-seq data, the scRNA-seq data (GSE166676) showed that myeloid cells (clusters 7, 8, 10, 16, 19, and 20) primarily synthesized *BAFF* (Figure S10A and B). Using immunohistochemistry, we further confirmed that BAFF staining colocalizes with CD11b+ (highly expressed by myeloid cells) in the AAA sections (Figure 6E and F). Interestingly, we did not find *APRIL* (a BAFF homolog)-expressing transcript clusters. Additionally, both scRNA-seq and Visium HD data revealed that B-cells, not T-cells, in the TLSs strongly expressed both *BR3 (TNFRSF13C)* and *TACI (TNFRSF13B)* (Figure S10A).

Altogether, these data suggest that myeloid BAFF is a possible modulator of B-cell autoreactivity during the development of AAAs in humans.

## DISCUSSION

The presence of both adipocytes and TLSs is strongly linked to aneurysm growth and rupture^13,21^. However, the roles of these vascular adipocytes and TLSs are poorly understood. Multiple reports have previously shown cross-sectional images of human AAAs in which adipocytes can be found close to TLSs^38,68,69^. However, the roles of these adipocytes were unclear. Here, we showed that TLSs were associated with adipocytes and, using multi-omics approaches, determined the putative roles of these vascular adipocytes and how they would support TLS formation by expressing factors that promote leukocyte migration and retention. Additionally, by comparing protein expression patterns in non-TLS lymphocyte aggregates in a control aorta and TLSs in a AAA biopsy, we identified unique intracellular pathways in TLSs, implicating them in TLS maturation.

PVAT is normally found on the abdominal aorta of healthy mice^70^ and is studied for its role in murine atherosclerosis^71,72^ and AAAs^73,74^. In contrast to mice, our study and images of human abdominal aorta cross-sections presented in previous reports^75–77^ suggest that PVAT is rudimentary in abdominal aortas of healthy humans. Adipocytes that appear in abdominal aortas of organ donors considered healthy may be due to aortic inflammation related to aging^78^, possibly analogous to intramuscular adipocytes formed in sarcopenia (which causes decreased muscle mass and strength), which is strongly linked to old age^79^. Our data showing adipocytes primarily in AAAs align with previous reports of increased adiposity in AAAs. One report indicates that the appearance of adipocytes in AAAs is due to adipogenic degradation of the aorta^21^. In human AAAs, expression of arterial wall-degrading factors and production of proinflammatory factors by adipocytes are positively correlated with AAA pathogenesis^23,80–82^. Indeed, microarray analyses of ruptured and non-ruptured human AAAs revealed a strong correlation of adipogenesis with ruptured AAAs^21^.

A critical question is: what are the precursor cells of these adipocytes? One report suggests mesenchymal stem cells as the origin of adipocytes^21^. A scRNA-seq study of human AAAs identified adipocyte-like SMCs, that is, SMCs expressing markers of adipocytes^19^. Therefore, we followed a rigorous approach to annotating adipocytes in our snRNA-seq experiment. Anatomically, adipocyte-TLS in AAAs looks similar to adipocyte-TLS in fatty lymph nodes (LN), in which CD34+ medullary stromal cells (fibroblasts) transdifferentiate into adipocytes, a process called LN lipomatosis^83^. Therefore, in human AAAs, we speculate that one of the four fibroblast subtypes identified in our snRNA-seq transdifferentiates into adipocytes. We also identified at least two ways adipocytes form in AAAs: one similar to the PVAT, and another in the medial-adventitial region. The latter adipocytes primarily localize with the TLSs. It is unknown what factors would trigger the generation of these adipocytes. One group created a hypoperfusion-induced murine model of AAA, which led to the generation of adipocytes, and the AAA histology looks similar to human AAAs with adipocytes^84^. This suggests a possible explanation for medial-adventitial adipocytes in AAA: increased hypoxia in the aortic tissue^85^, possibly due to the expansion of the aorta and constriction of vasa vasorum.

LN lipomatosis is associated with aging^86^ and cancer patients^87^, and is thought to be one of the mechanisms of impaired immunity to infections and vaccines^88^. Therefore, we investigated whether the vascular adipocytes express molecules that affect TLS formation. Our snRNA-seq data revealed that vascular adipocytes produce various factors that attract leukocytes and provide support for immature TLS formation. A previous study on the *ex vivo* culture of adventitial and medial explants from AAA tissues demonstrated that both explants release chemokines involved in TLS formation, such as CCL19, CCL20, CCL21, CXCL13, and CXCL16^13^. Our spatial transcriptomics data localized these chemokines to TLSs. A previous immunohistochemical staining of AAA sections^13^ revealed that these chemokines colocalized with cells expressing aSMA, suggesting the possibility of SMCs as TLSo. In this line, we also identified SMCs in TLSs. However, SMCs as TLSo need further validation, as aSMA is also expressed by fibroblasts in the medulla and paracortex of lymph nodes and promotes SLO formation^83^. Proliferating CD21+CD44+ stromal cells^68^ and CXCL12+ fibroblasts^89^ may also function as TLSo. In this line, our study provides crucial information that vascular adipocytes, by synthesizing various factors, can also act as non-canonical TLSo, particularly those localized in the medial-adventitial region. For example, thrombospondin 1 can be both a chemotactic factor by attracting leukocytes, possibly from the vasa vasorum, and a hapatotactic factor that promotes binding of leukocytes to the adipocyte surface^90^. Autotaxin is one of the chemotactic factors that can increase the levels of bioactive proinflammatory lipid lysophosphatidic acid in damaged AAA tissues^26^. Additionally, we found that some adipocytes lack the crucial droplet-binding enzyme Plin1, which renders adipocytes dysfunctional^36^. We recently reported that matrix metalloproteinase-2 (MMP2) impairs the activity of glucose transporter 4 (GLUT4) on the surface of adipocytes and negatively impacts adipocyte glycolysis^91^, a critical metabolic pathway for adipocyte function. Because active MMP2 concentrations are high in AAA tissues^92^, it is possible that MMP2 creates dysfunctional adipocytes by attenuating glycolysis, leading to reduced Plin1 expression. Thus, vascular adipocytes, unexpectedly, have the potential to serve as non-canonical TLSo in human AAAs.

Formation of TLSs and generation of autoantibodies in human AAAs was first described by Tilson et al.^93,94^. Flow cytometry studies performed on human AAA tissues revealed the presence of CD19+ B cells, CD8+ T cells, CD4+ T cells, and CD4+CD45RO+ memory T cells^13^. Our multiomics study revealed that TLSs within AAAs recapitulate the ontogeny of SLOs. A previous study demonstrated that adventitial tissues from AAAs, but not from control aortas, produce higher levels of IgGs^13^. However, it was unclear whether TLS-derived plasma cells or antigen-experienced plasma cells infiltrating from the vasa vasorum synthesized the IgGs. Our Visium data clearly demonstrates immunoglobulin transcripts for IgG1, IgG3, IgA, and IgM in TLSs. This result, along with the presence of secondary B-cell follicles, suggests that the TLSs within AAAs produce antibodies. In murine models of AAA, such as the topical elastase model, clusters of B-cells and T-cells were observed^7^, which may represent immature TLSs. However, aged apolipoprotein E–deficient (ApoE-/-) mice with advanced atherosclerosis exhibit mature TLS organization localized near plaques^95,96^. Therefore, AAA studies on aged ApoE-/-mice may mimic advanced human AAAs and be helpful to determine the extent to which the autoantibodies targeting the aorta are pathogenic and exacerbate AAA growth.

Our screening for IgG autoantibodies confirmed a few of the previously reported AAA-associated autoAbs, such as anti-collagen and anti-elastin^97^. Of interest, culture of AAA explants identified anti-EBNA1 Ab as a dominant secreted autoAb. Epstein–Barr virus infection and the presence of anti-EBNA1 Ab are identified in over 95% of the US population aged ≥40^55^. A small percentage of infected individuals develop cancer or autoimmune diseases in their lifetime. EBNA1 is known to share conformational and amino acid sequence similarities with human proteins such as Myelin Basic Protein, Aquaporin-4, and GliaCAM^98^, and therefore, anti-EBNA1 Abs cross-react with those proteins. In agreement with autoreactive B-cells producing autoAbs, clonally expanded αβ T-cell receptor+ T-cells were identified in AAA tissues^99^. In addition, autoAbs against intracellular components such as CENP-B suggest that progressive tissue damage provides a continuous supply of autoantigens and that an abnormal local immune response uses these autoantigens to promote inflammation. Related to B cell-mediated vascular immunity, a recent article demonstrates the generation of high-affinity pathogenic anti-histone antibodies in the TLSs of murine atherosclerotic lesions^100^.

BAFF exists as 3-mer and 60-mer forms and differentially activates canonical and non-canonical NF-kB signaling^67^. In particular, the 60-mer is the highly active form of BAFF and can transcriptionally and metabolically hyperactivate B cells. On the other hand, BCR crosslinking by antigens phosphorylates CD79a and CD79b and activates Syk-dependent signaling^101^. In an inflammatory tissue milieu, such as AAA, B cells are highly likely to be exposed to both BAFF and antigens. Schweighoffer *et al.* have reported that BAFF signaling *via* the BR3 uses the phosphorylated CD79a and Syk (generated by BCR activation) as adaptor proteins, to transduce survival signals *via* ERK1/2 phosphorylation^66^. Our NanoString Spatial Protein Profiling and immunohistochemistry together revealed that ERK1/2 and canonical NF-kB pathways are activated in B cells localized in non-TLS aggregates within a BAFF microenvironment. This suggests the possibility that these B cells respond to BAFF and local antigens, and BAFF 60-mer promoted the survival of autoantigen-exposed B cells. However, because of technical limitations, we cannot determine whether BAFF in non-TLS and TLS environments exists as 3-mers or 60-mers.

Our study has several limitations. Our current interpretation of TLS anatomy is based on multiple 5 µm cross-sections of a large piece of AAA tissue, at least >4 cm2 in area and 0.3-0.5 cm thick. Although we can identify TLSs associated with adipocytes, we may have missed some associations because adipocytes may present in other cross-sections along the thickness. Similarly, the absence of TLS and adipocytes in a tissue cross-section does not indicate that TLS and adipocytes are absent in the entire AAA tissue. In this line, the advanced TLSs in AAA13 had adipocytes in early sections (Figure 3A), but not in the later sections (Figure 5B). Another limitation is the number of frozen biospecimens used for our snRNA-seq and VisiumHD transcriptomics studies. Our rationale for selecting 4 specimens was based on histopathological analysis of FFPE sections from another part of the same AAA for the presence of vascular adipocytes and TLSs. However, inflammation may differ between the tissue used for histopathology and the tissue used for snRNA-seq. Therefore, we performed immunohistochemical analyses to confirm protein-level expression of genes of interest identified in snRNA-seq studies. Additionally, for nuclei preparation for the snRNA-seq, we used two 100 mg tissue pieces/sample. These two tissue pieces are unlikely to represent the entire AAA pathology. In our VisiumHD studies, we largely missed gene expression signatures from adipocytes because they have a very thin layer of cytoplasm and an eccentric nucleus. Lastly, more studies are needed to determine if human AAA is an autoimmune-driven disease or if autoimmunity exacerbates existing vascular degeneration and inflammation.

Overall, we described a possible role for vascular adipocytes in organizing leukocyte cluster formation, which then matures into TLS and promotes B cell-mediated inflammation within aneurysm tissue. While more studies are needed to confirm these findings, this work opens new avenues of investigation for potential therapeutics for treating small AAAs. Particularly, targeting adipogenesis, possibly with anti-diabetic drugs, may attenuate B-cell inflammation and AAA growth. Additionally, anti-BAFF therapeutics could neutralize BAFF in TLSs, allowing autoreactive B cells to die and mitigating inflammation in AAAs.

## DISCLOSURE

None

## ACKNOWLEDGEMENTS

We thank Jerald Whitley for technical assistance, and the East Carolina University Research Histology Core and Imaging Core for technical support. The illustration in the graphical abstract was created with BioRender.com.

## FUNDING

This work was supported by NIH R01 HL146685, Brody Brothers Endowment Award, and funds from East Carolina University to AM.

## Nonstandard Abbreviations and Acronyms

AAA: Abdominal Aortic Aneurysm
APRIL: A Proliferation-Inducing Ligand
AutoAb: Autoantibody
BAFF: B-cell-activating factor
BCMA: B-Cell Maturation Antigen
BR3: BAFF Receptor / BAFF-R
FFPE: Formalin-Fixed Paraffin-Embedded
Ig: Immunoglobulin
KEGG: Kyoto Encyclopedia of Genes and Genomes
PVAT: Perivascular adipose tissue
SMC: Smooth muscle cell
SLE: Systemic Lupus Erythematosus
scRNA-seq: Single-cell RNA sequencing
snRNA-seq: Single-nucleus RNA sequencing
TACI: Transmembrane Activator and CAML Interactor
TNF: Tumor Necrosis Factor
TLS: Tertiary lymphoid structures
TLSi: TLS inducers
TLSo: TLS organizing cells

## SUPPLEMENTAL FIGURE

**Figure S1.**
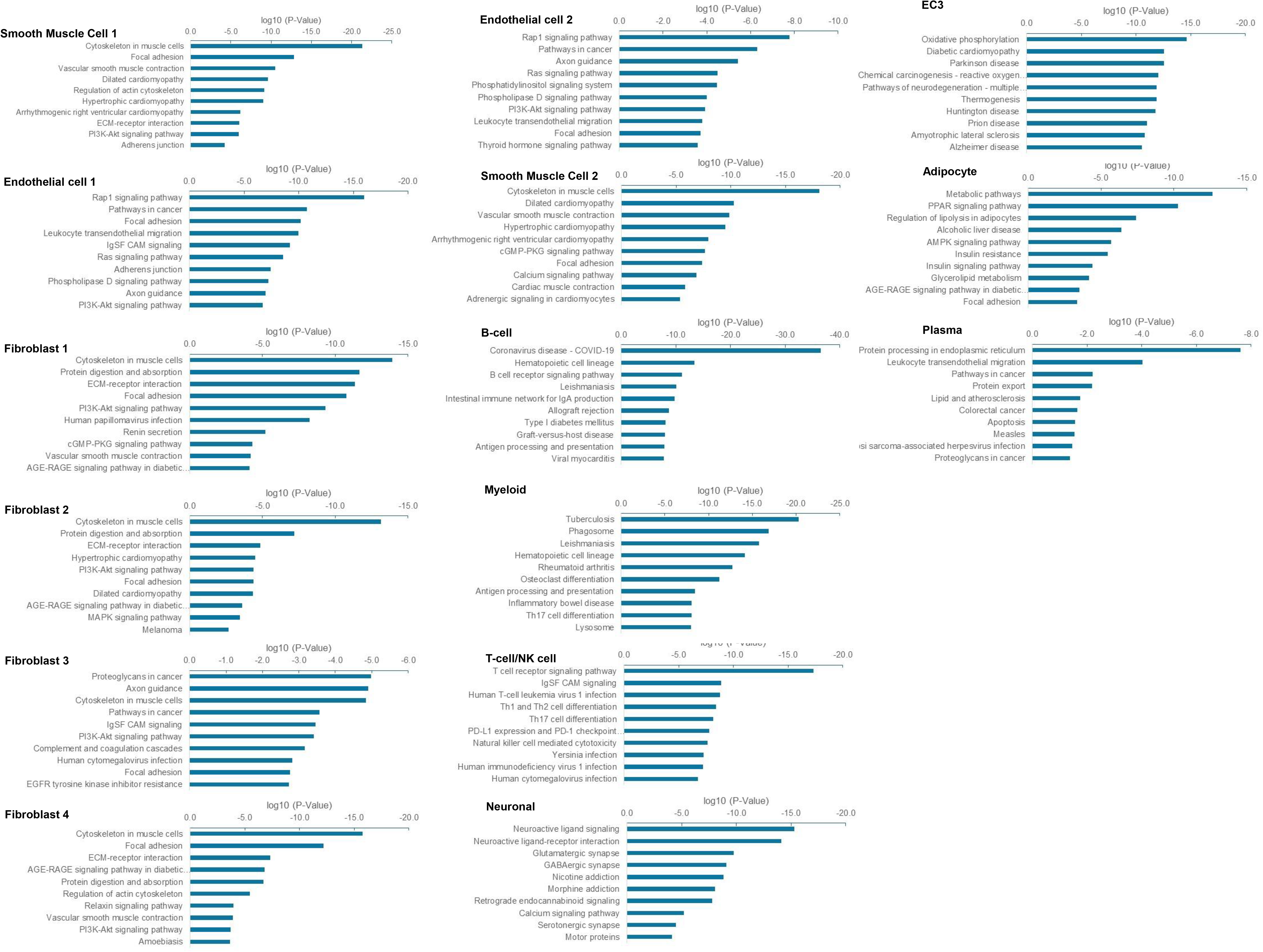
KEGG (Kyoto Encyclopedia of Genes and Genomes) pathway analysis of clusters identified in snRNA-seq of human AAAs. Highly expressed genes (adjusted p-value <0.05) from each cluster in the snRNA-seq data were used for KEGG pathway analysis, and the top ten statistically significant (based on modified Fisher’s exact test p-values) biological pathways are shown.

**Figure S2.**
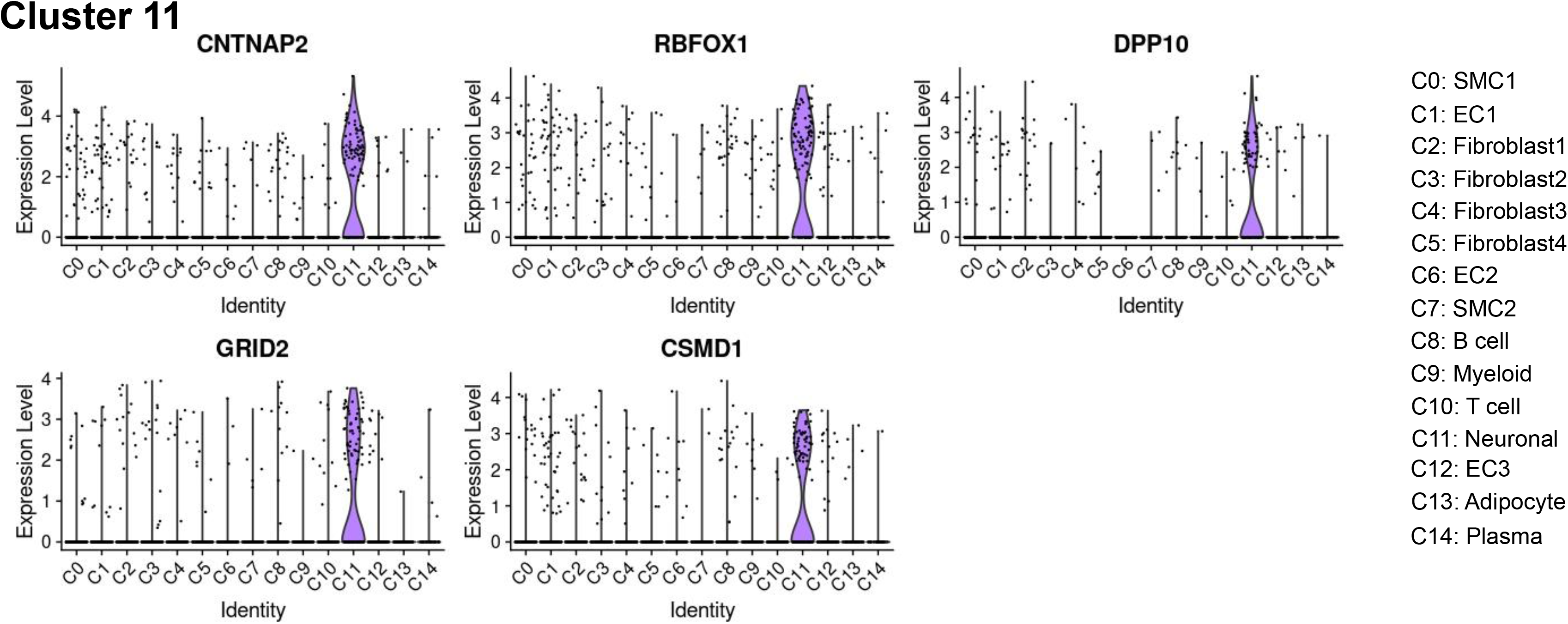
In silico analysis of cluster 11 in the snRNA-seq data of human AAAs. Violin plots showing comparative expressions of genes highly expressed in cluster 11.

**Figure S3.**
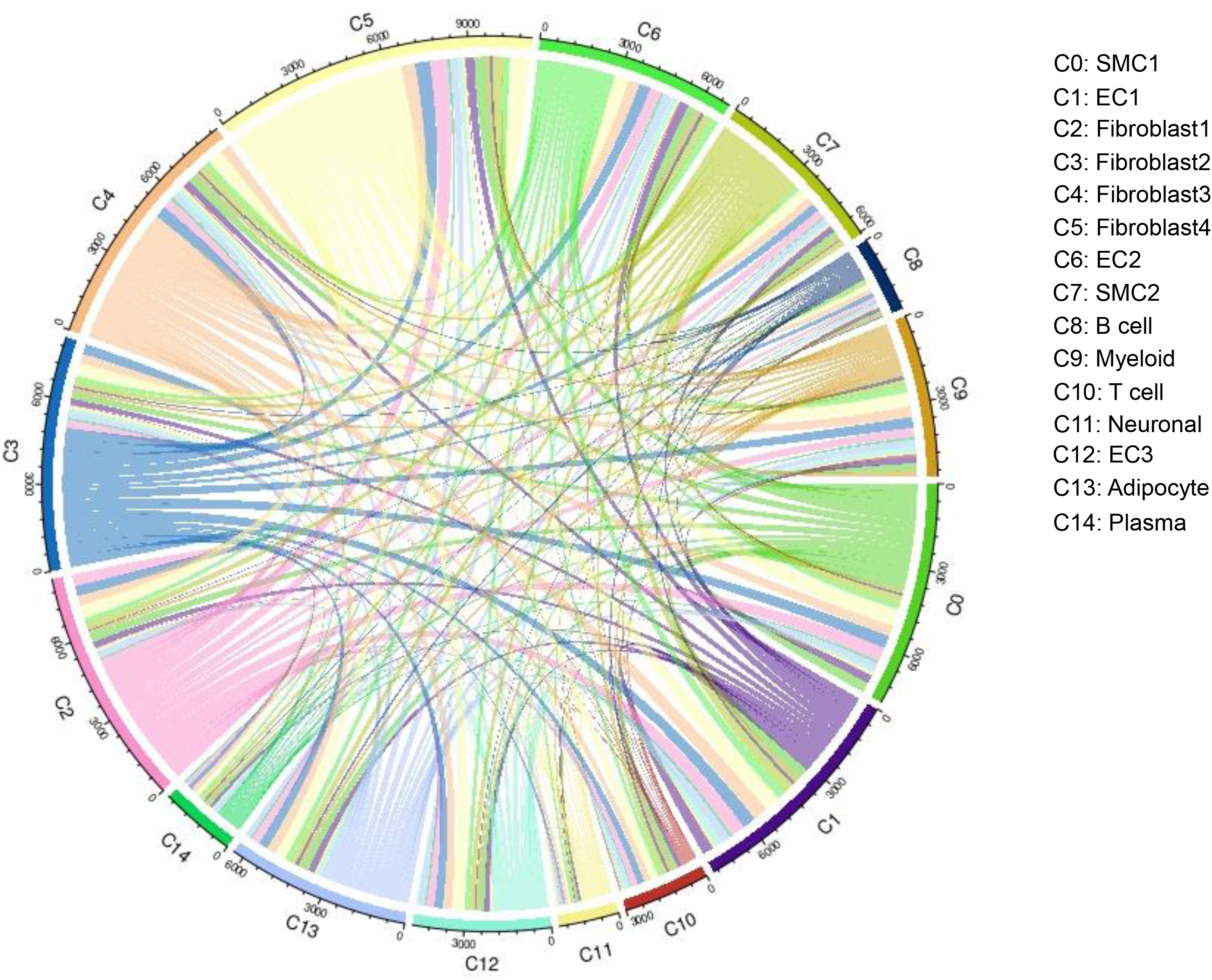
scDiffCom analysis of the probability of cell-cell interaction among cellular groups identified by snRNA-seq. Chord diagram shows the probability of active interactions among the 14 cell groups identified in the human snRNA-seq of human AAAs.

**Figure S4.**
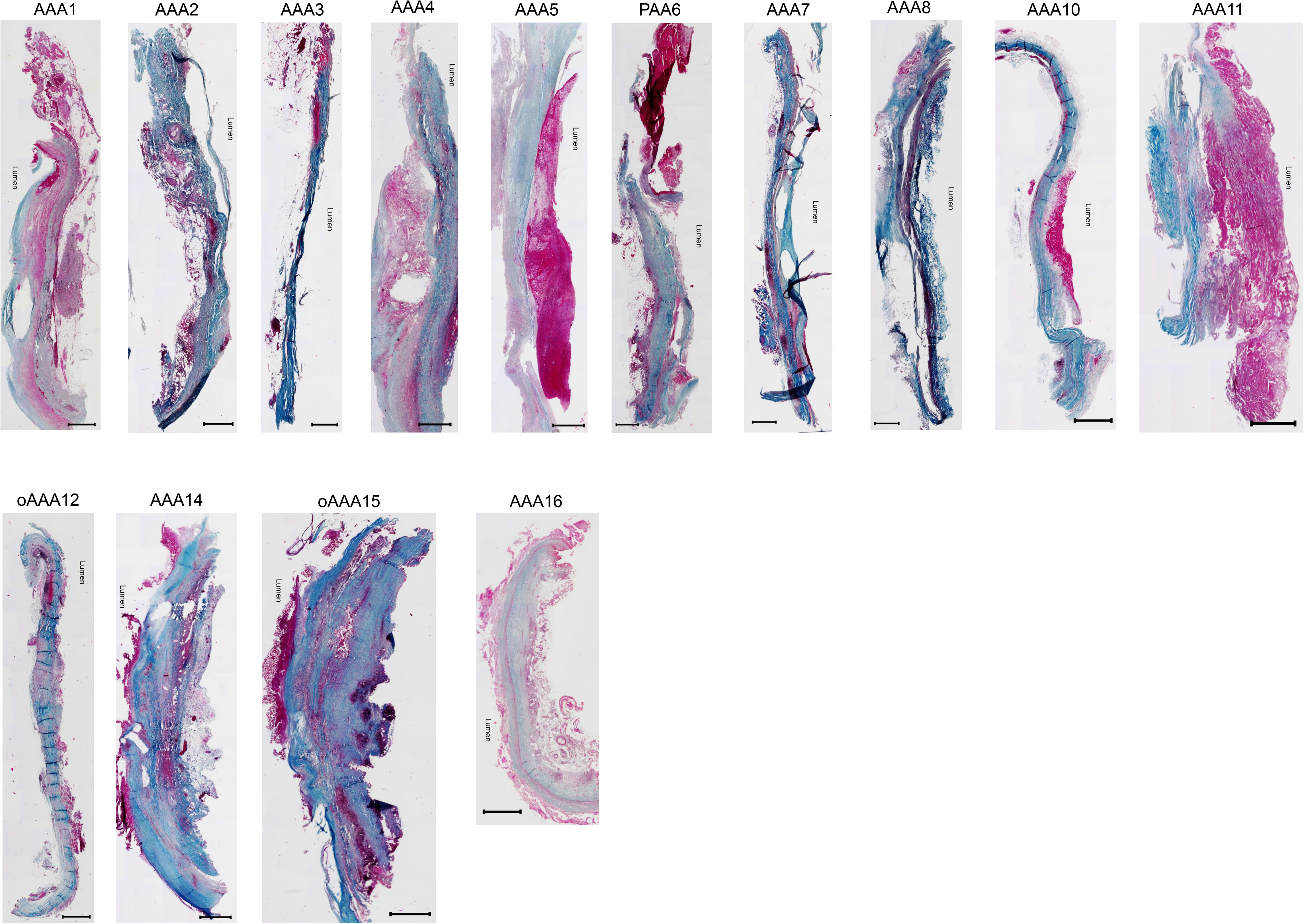
Histopathology of AAAs. Whole-slide images of Masson’s trichrome staining of cross-sections from 13 AAA patients and 1 PAA patient. Blue: collagen; red: muscle and cytoplasm; black: nuclei. Scale bars are shown in each image. Scale bars are shown in each image.

**Figure S5.**
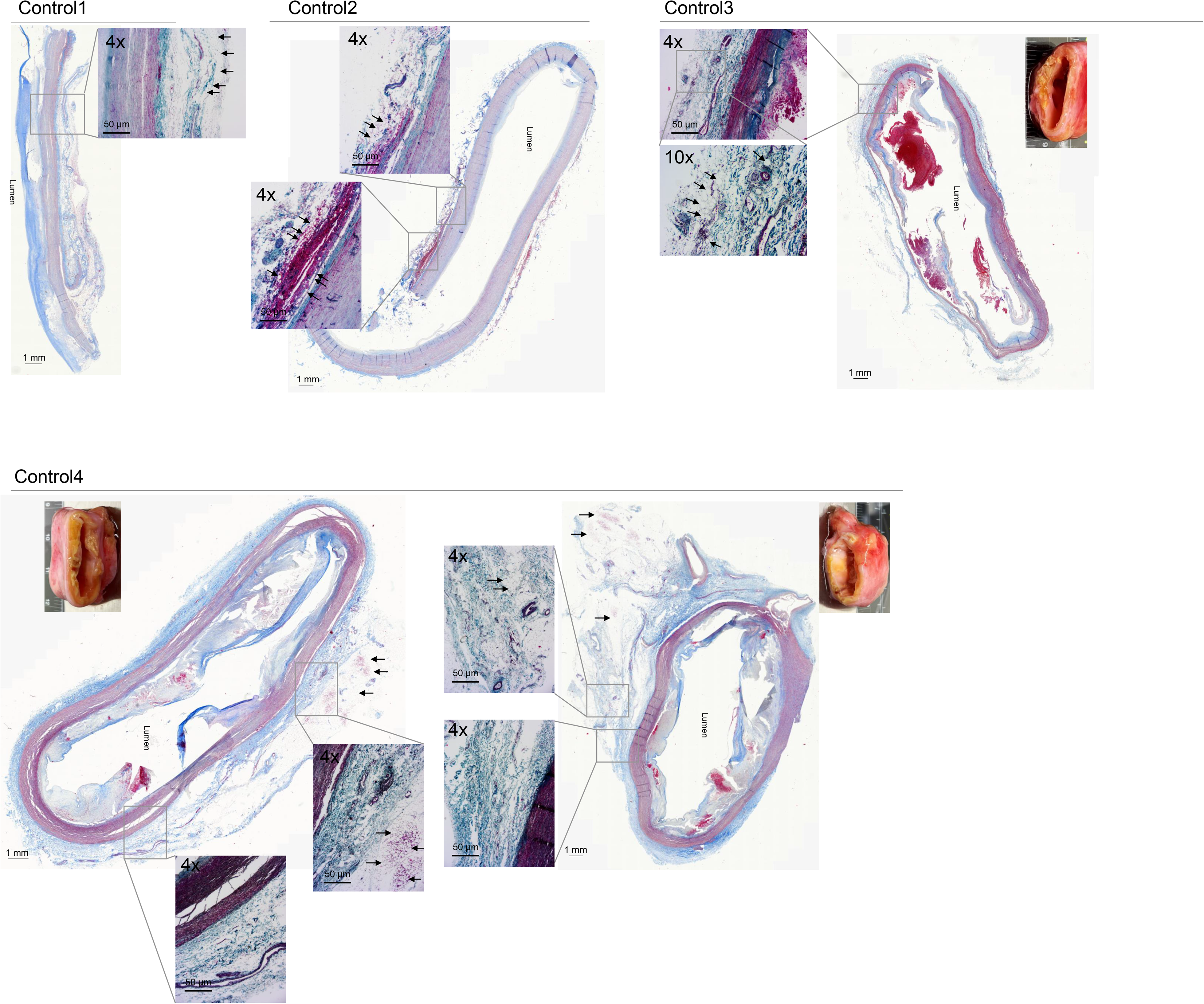
Histopathology of control abdominal aortas and localization of vascular adipocytes. Whole-slide images of Masson’s trichrome staining of cross-sections from 4 control abdominal aortas (acquired using MoticEasyScan), with single-field images (acquired using Laxco LMI) of the region of interest and unilocular white adipocytes identified by arrows. The inset in the stained aortas shows unmanipulated tissue used for paraffin embedding, followed by cross-sectioning.

**Figure S6.**
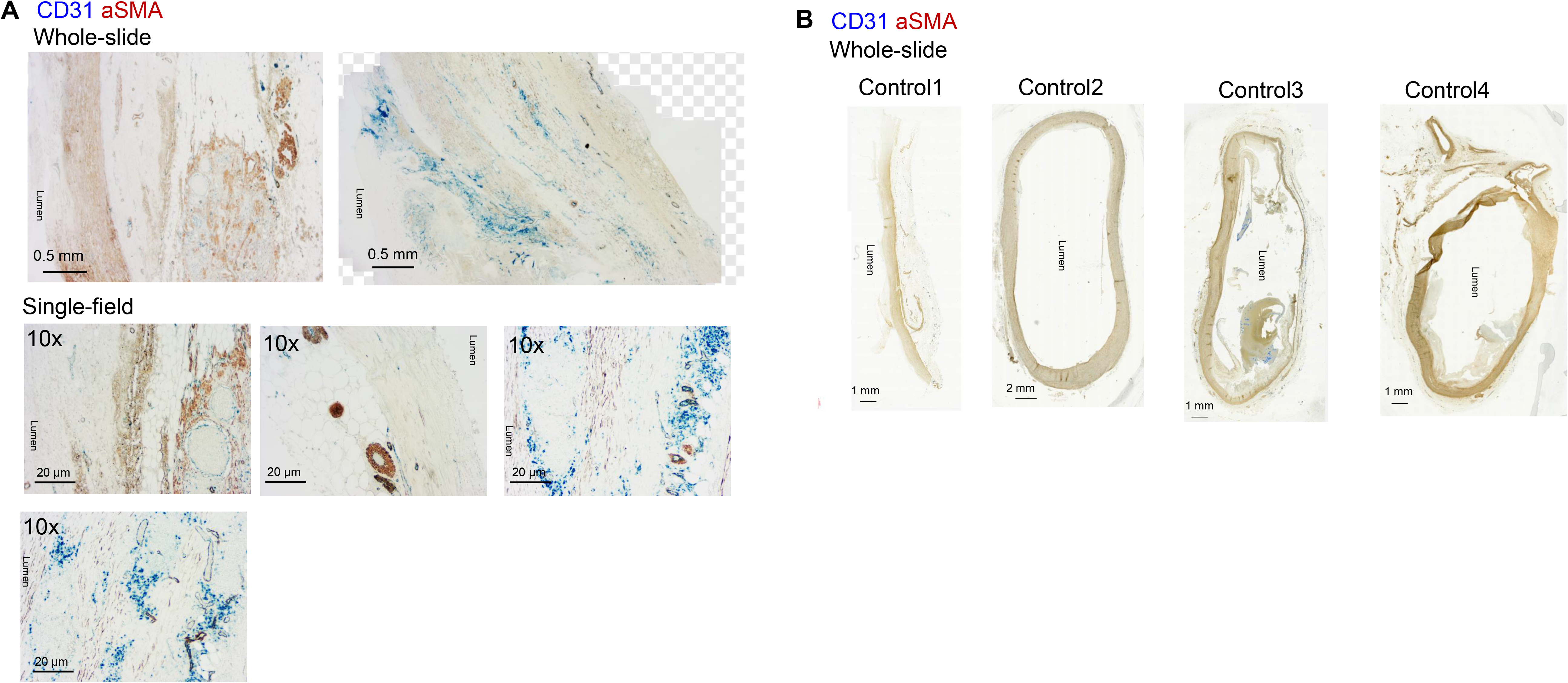
Endothelial and smooth muscle cell layers in AAA and control abdominal aortas. A, Whole-slide and single-field images of AAA cross-sections showing endothelial cells (CD31, blue) and smooth muscle cells (aSMA, brown). **B,** Whole-slide images of non-aneurysmal control abdominal aortas showing endothelial cells (CD31, blue) and smooth muscle cells (aSMA, brown). Scale bars are shown in each image; n=6 patients and 4 control abdominal aortas.

**Figure S7.**
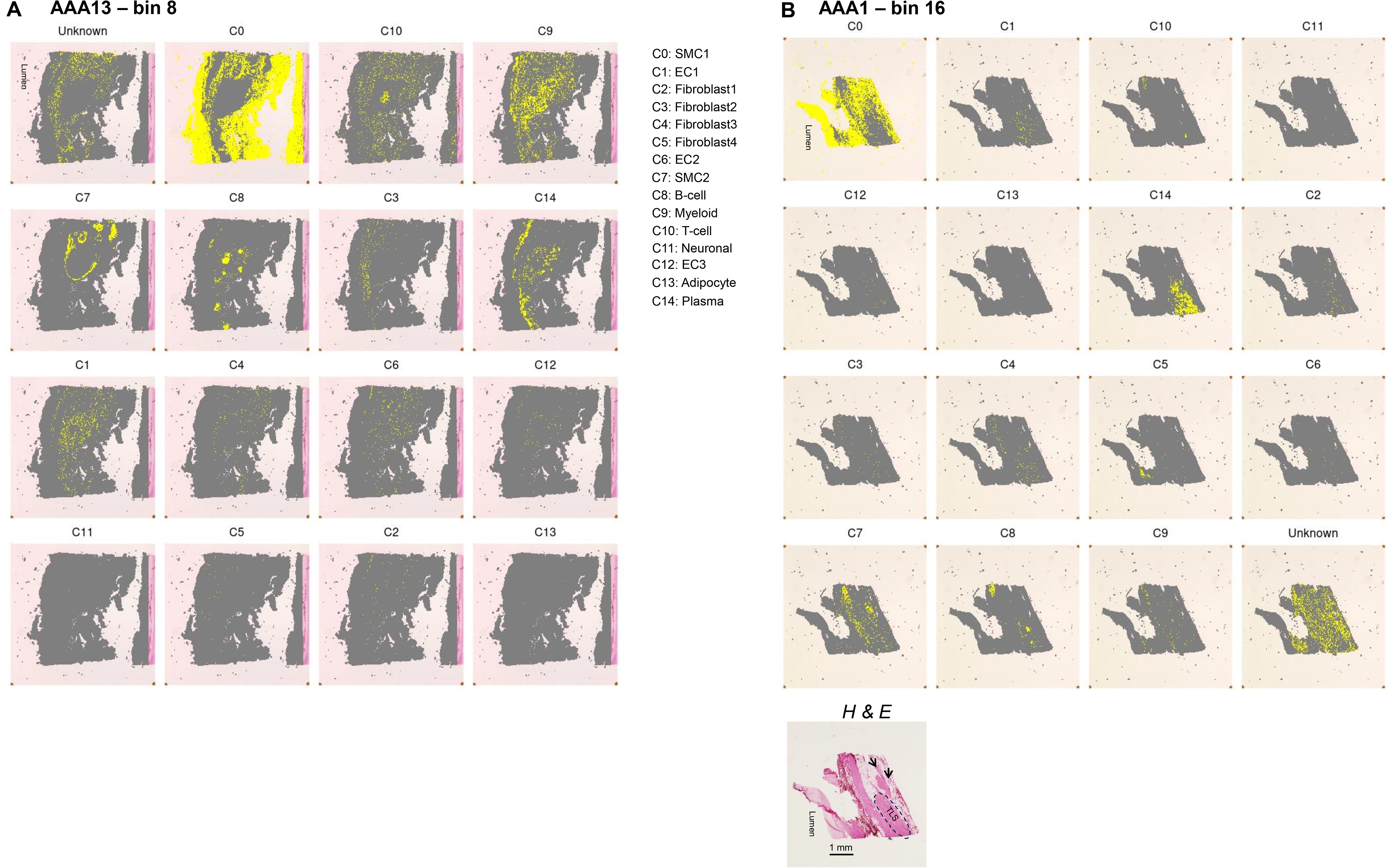
Visium HD spatial transcriptomics on human AAA13 and AAA1. **A**, 8 µm (8 x 8) bins showing localization of all 14 cellular clusters obtained from snRNA-seq on the AAA13 cross-section. B, 16 µm (8 x 8) bins showing localization of all 14 cellular clusters obtained from snRNA-seq on the AAA1 cross-section. Hematoxylin and Eosin (H & E)-stained image of AAA1 used for Visium HD spatial transcriptomics; scale bar = 1 mm.

**Figure S8.**
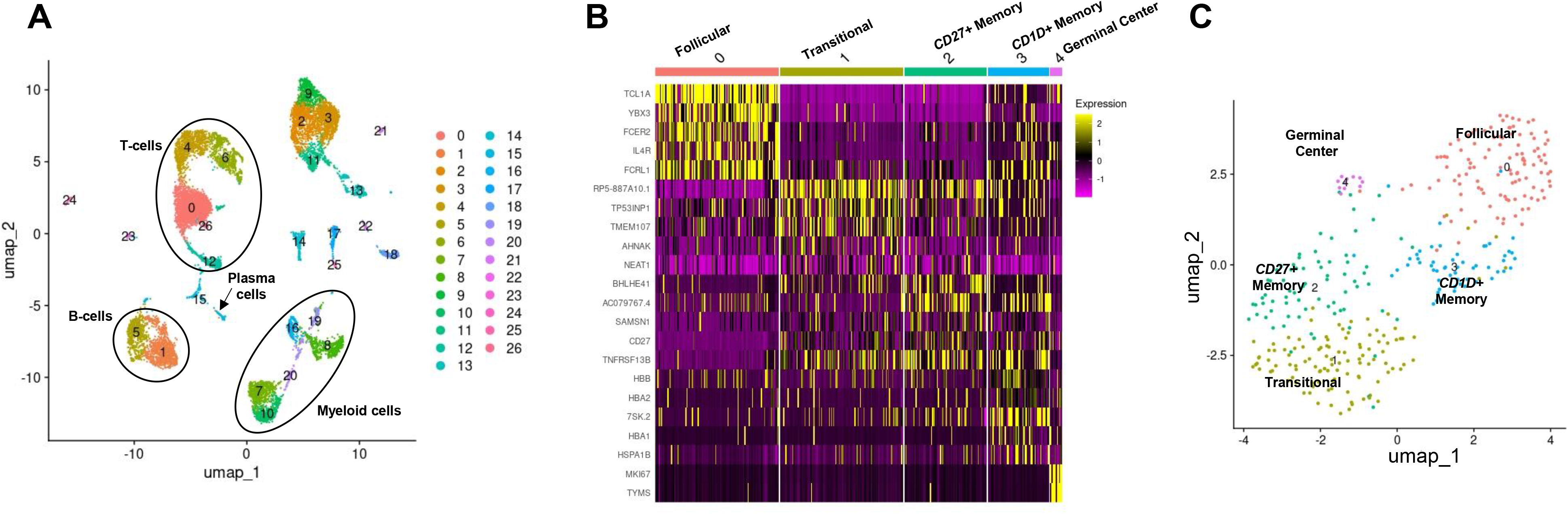
Re-analysis of scRNA-seq human AAA data GSE166676 for B-cells. **A**, Uniform Manifold Approximation and Projection (UMAP) plot showing 26 cellular clusters from scRNA-seq of AAA specimens from 3 patients and control aortic tissues from 2 individuals. B cells, plasma cells, T cells, and myeloid cells are indicated. **B**, Heat plot illustrating the top five marker genes for each B-cell cluster shown in the UMAP plot in **C**.

**Figure S9.**
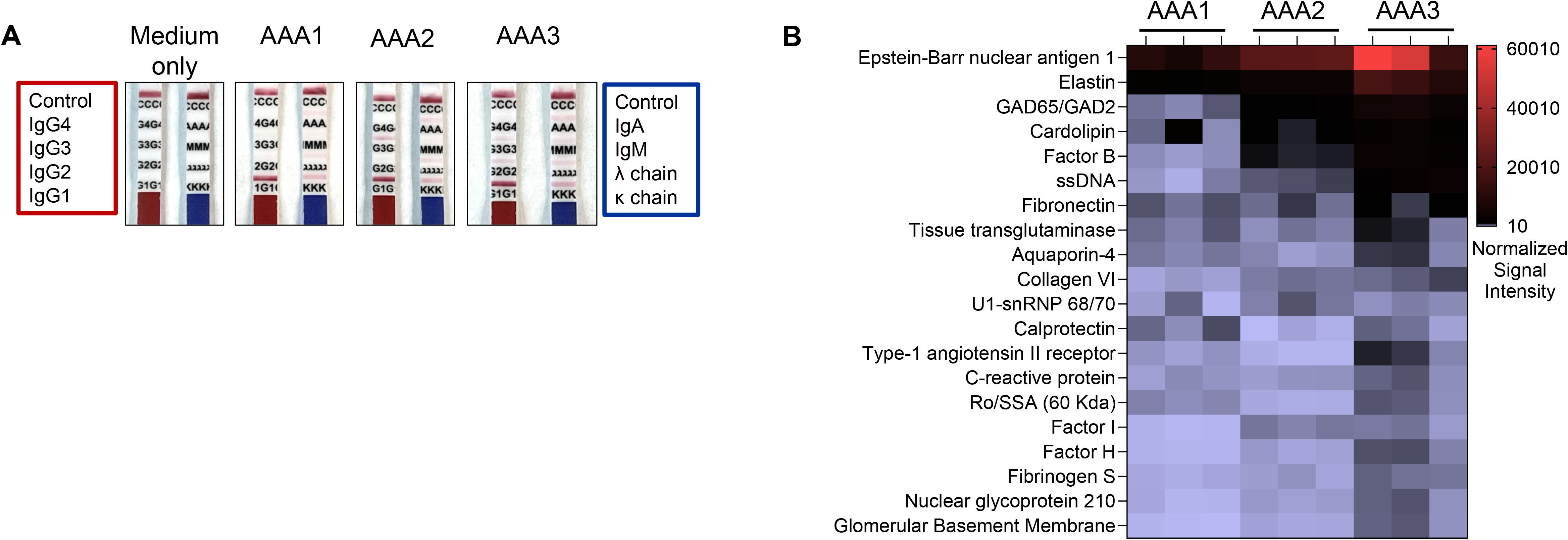
Qualitative assessment of immunoglobulins secreted by AAA explants. **A**, Culture supernatants from AAA1, AAA2, and AAA3 explants (n=3 explants/AAA tissue) were examined for the presence of immunoglobulin isotypes using test strips. Representative images are shown. **B**, Quantification of the top 20 IgG autoantibodies in the culture supernatants of AAA1, AAA2, and AAA3 explants (n=3 explants/AAA tissue). Signal intensity was normalized to the culture medium without explant.

**Figure S10.**
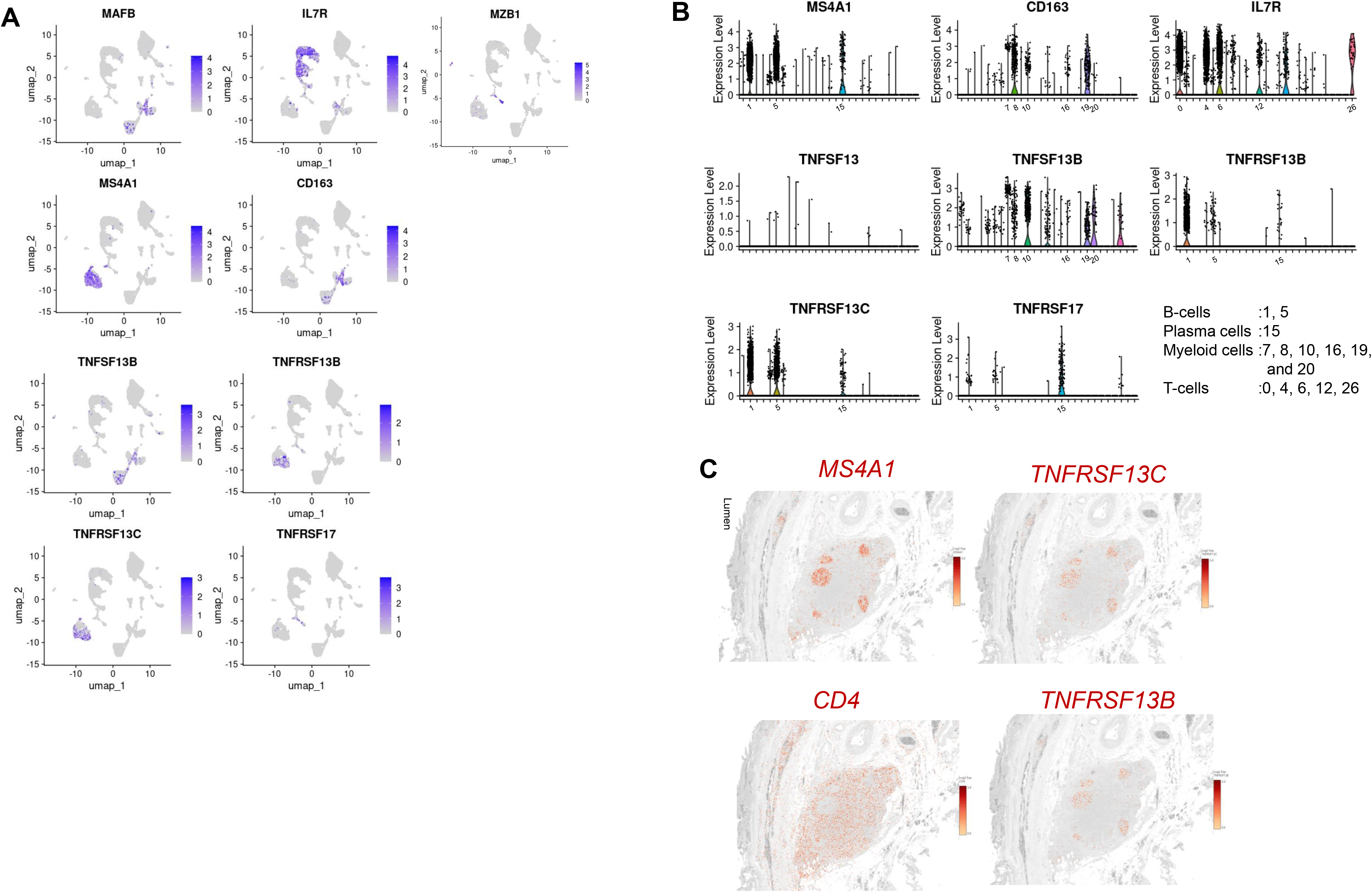
ScRNA-seq human AAA data GSE166676 revealed that B cells primarily express the BAFF receptors BR3 and TACI. **A**, UMAP plots illustrating clusters highly expressing myeloid cell markers *MAFB* and *CD163*, T-cell marker *IL7R*, and B-cell marker *MS4A1*, plasma cell marker *MZB1*, followed by BAFF (*TNFSF13B*), TACI (*TNFRSF13B*), BR3 (*TNFRSF13C*), and BCMA (*TNFRSF17*). **B**, Violin plots illustrate comparative gene expression of B cells, myeloid cells, and T-cells, followed by APRIL (TNFSF13), BAFF (TNFSF13 B), and BAFF receptors TACI, BR3, and BCMA. **C**, 16 µm bins of Visium HD spatial transcriptomics on AAA13 showing spatial images with gene expression patterns to illustrate areas of B-cells (*CD79A1*) and CD4+ T cell (*CD4*), followed by TACI (*TNFRSF13B*) and BR3 (*TNFRSF13C*). Gene expression intensities are displayed.

## Notes

### Competing Interest Statement

The authors have declared no competing interest.

### Summary of Updates

New title, Abstract was revised, minor changes in the text, a part of the Figure 5 was moved to Figure S9, issues with Table S2 fixed, and a BioRender acknowledgment statement was added.

