## Supplementary material for "Spatial multi-omics implicates vascular adipocytes proximal to tertiary lymphoid structures in human aortic aneurysm and B-cell autoreactivity": Table S2

**Supplemental Table 2**

**Functionality of highly expressed genes in vascular adipocytes**

| Gene ID | Gene name | Function |
|---|---|---|
| LPL | Lipoprotein lipase | Catalyzes the hydrolysis of triglycerides, plays an important role in lipid clearance from the bloodstream, lipid utilization, and storage. |
| PLIN5 | Perilipin 5 | Recruits mitochondria to the surface of lipid droplets and is involved in lipid droplet homeostasis by regulating both the storage of fatty acids in the form of triglycerides and the release of fatty acids for mitochondrial fatty acid oxidation. |
| PLIN1 | Perilipin 1 | Coats lipid storage droplets to protect them from breakdown by hormone-sensitive lipase |
| CIDEA | Cell death inducing DFFA like effector A | Lipid transferase that promotes unilocular lipid droplet formation by mediating lipid droplet fusion |
| ADIPOQ | Adiponectin | Important adipokine involved in the control of fat metabolism and insulin sensitivity, with direct anti-diabetic, anti-atherogenic and anti-inflammatory activities |
| MLXIPL | MLX interacting protein like | This gene encodes a basic helix-loop-helix leucine zipper transcription factor of the Myc/Max/Mad superfamily. This protein forms a heterodimeric complex and binds and activates, in a glucose-dependent manner, carbohydrate response element motifs in the promoters of triglyceride synthesis genes |
| THBS1 | Thrombospondin-1 | Secreted adhesive glycoprotein that mediates cell-to-cell and cell-to-matrix interactions, regulates platelet aggregation, angiogenesis, and tumorigenesis |
| ENPP2 | Ectonucleotide pyrophosphatase/phosphodiesterase 2 (Autotaxin) | Secreted lysophospholipase D that hydrolyzes lysophospholipids to produce the signaling molecule lysophosphatidic acid in extracellular fluids (lysophosphatidic acid regulates cell proliferation, survival, migration, angiogenesis, and immune responses) |
| AQP7 | Aquaporin 7 | Aquaglyceroporins form homotetrameric transmembrane channels, with each monomer independently mediating glycerol and water transport across the plasma membrane along their osmotic gradient, regulates lipolysis and glycerol efflux in adipocytes |
| SLC7A10 | Solute carrier family 7 member 10 | Associates with SLC3A2/4F2hc to form a functional heterodimeric complex that translocates small neutral L- and D-amino acids across the plasma membrane, regulates the fates of white and beige/brown adipocytes. |
| PDE3B | Phosphodiesterase 3B | Cyclic nucleotide phosphodiesterase that hydrolyses the second messengers cAMP and cGMP, and regulates key intracellular signaling pathways and many important physiological |

| | | |
|---|---|---|
| | | processes. Involved in negative regulation of angiogenesis, negative regulation of cell adhesion, and negative regulation of lipid catabolic processes. |
| GHR | Growth hormone receptor | This gene encodes a member of the type I cytokine receptor family, which is a transmembrane receptor for growth hormone. Binding of growth hormone to the receptor leads to receptor dimerization and the activation of an intra- and intercellular signal transduction pathway, leading to growth |
