## Supplementary material for "Spatial multi-omics implicates vascular adipocytes proximal to tertiary lymphoid structures in human aortic aneurysm and B-cell autoreactivity": Table S4

**Supplemental Table 2**

**Aortic aneurysm specimens**

| Sl. No. | Disease | Age and sex | Medical conditions | Drugs | Vascular adipocytes | TLS | IVA with TLS |
|---|---|---|---|---|---|---|---|
| 1 | AAA1 | 65y, male | Obesity, smoker | N/A | Yes | Yes | Yes |
| 2 | AAA2 | 70y, male | Obesity, smoking, Coronary Artery Disease, atrial fibrillation | N/A | Yes | Yes | Yes |
| 3 | AAA3 | 70y, male | Diabetes, hypercholesterolemia, peripheral vascular occlusive disease, hypertension, and tobacco abuse | N/A | Yes | Yes | Yes |
| 4 | AAA4 | 61y, female | PMH: hypertension, gerd, benign colon neoplasm, smoker, PAD | Pletal, Lipitor, Hctz, lisinpril | Yes | Yes | Yes |
| 5 | AAA5 | 73y, female | chronic obstructive pulmonary disease, hypertension | Metoprolol, ampicillin, cilostazol | No | No | No |
| 6 | AAA7 | 80, male | Polysubstance abuse, hypertension, Diabetes, tobacco abuse | Amlodipine, aspirin, atorvastatin, vitamin D3, donepezil, doxazosin, finasteride, gabapentin, glipizide, lisinopril, metoprolol, spironolactone | Yes | Yes | No |
| 7 | AAA8 | 61y, male | Gastroesophageal reflux disease, Diabetes, hepatitis C, hypertension, Hyperlipidemia | Metformin, naproxen, omeprazole, amlodipine, atorvastatin, aspirin | Yes | No | No |
| 8 | AAA10 | 68y, male | Hodgkin's lymphoma, Diabetes, stroke | Tamsulosin, Xarelto, pyridium, metformin, melatonin, evolucumab, cyclobenzprine, aspirin | Yes | Yes | No |
| 9 | AAA11 | 66, male | Alcoholism, Hypertension, coronary artery disease, myocardial infarction, hypothyroidism, hyperlipidemia, diabetes, Smokes cigarettes daily | Testosterone, pravastatin, protonix, metoprolol, lisinopril, levothyroxine, hydrochlorothiazide, clopidogrel, aspirin, baclofen, famotidine | No | No | No |

| | | | | | | | |
|---|---|---|---|---|---|---|---|
| 10 | AAA12 (Occlusive) | 79, male | Chronic obstructive pulmonary disease, hypertension, peripheral vascular occlusive disease | Beta-blockade, calcium-channel blocker, ciprofloxacin | Yes | Yes | Yes |
| 11 | AAA13 | 54y, male | hypertension and chronic back pain | No | Yes | Yes | Yes |
| 12 | AAA14 | 79, female | pulmonary embolism, Gastroesophageal reflux disease, hypertension, migraines | trileptak, Prozac, creon | Yes | Yes | Yes |
| 13 | AAA.15 (Occlusive) | 65, male | asthma, cirrhosis, peripheral vascular occlusive disease | No | Yes | Yes | Yes |
| 14 | AAA16 OriGene | 66, male | N/A | N/A | Yes | Yes | Yes |
| 15 | PAA6 Popliteal artery aneurysm | 63, male | Lyme disease | Aspirin, rosuvastatin, famotidine, eliquis | Yes | Yes | Yes |

**Control aortic specimens**

| Sl. No. | Disease | Age and sex | Medical conditions | Drugs* | Vascular adipocytes | TLS | IVA with TLS |
|---|---|---|---|---|---|---|---|
| 1 | Control 1 | 86, female | Acute pulmonary insufficiency, Acute blood loss anemia, Anxiety, Depression, NSTEMI, Falls, GERD, Subdural hematoma, Acute respiratory failure, Drug withdrawal hypersomnia, Mitral regurgitation, Tricuspid regurgitation, Hysterectomy, Closed fracture of neck of right femur, Mild pulmonary hypertension | Plavix, low-dose aspirin, Effexor, eye drops, heart medicine, Tylenol, Amiodarone, Aspirin, Vitamin D3, Cefuroxime, Clopidogrel, Cyanocobalamine, Loperamide, Melatonin, Omeprazole, Ondansetron, Calcium, Coenzyme | Yes | No | No |
| 2 | Control 2 | 70, female | Gastric Antral Vascular Ectasia (GAVE) syndrome, Alcohol use disorder (AUD - in remission for 10 years), Osteoporosis, Hepatic | Nortriptyline, duloxetine, Orencia injection, hydroxychloroquine, vitamin D, Iron, Vitamin B12 gummies, Tylenol Extra Strength, | A few clusters | No | No |

| | | | | | | | |
|---|---|---|---|---|---|---|---|
| | | | encephalopathy & Gastric varices, Chronic Iron deficiency anemia (IDA), Decompensated cirrhosis, Rheumatoid Arthritis (RA - previously on Abatacept & currently on Hydroxychloroquine), Ascites, Neutropenia, Microcytic anemia, Resp alkalosis, Peripheral Neuropathy, Gastric varices, Major Depressive Disorder (MDD), Anxiety | Tylenol pm, steroids, Lactulose, Lidocaine, Lorazepam, Nortriptyline, Omeprazole-Sodium bicarbonate, Spironolactone | | | |
| 3 | Control 3 | 66, male | Pancreatic metastatic cancer, Hyperlipidemia, Candidiasis, Transaminitis' Gastroesophageal reflux disease' Hyponatremia, Pulmonary embolism' Pneumonia, Pleural effusion, Prostate Cancer | Creon, Metoclopramide, Ondansetron, Mirtazapine, Omeprazole, Albuterol, Vitamin D, Multi-Vitamin | A few clusters | No | No |
| 4 | Control 4 | 63, female | Cerebrovascular accident, Chronic obstructive pulmonary disease, Depression | Morphine, Versed, Potassium Chloride (KCl), Potassium Bicarbonate, Duoneb, Vancomycin, Pepcid, Zosyn, Peridex, Keppra, Chlorhexidine, Levetiracetam, Midazolam, Ondansetron, Ciprofloxacin, Glucose (Dextrose), Famotidine, Ipratropium Bromide and Albuterol Sulfate, Piperacillin and Tazobactam, Insulin | A few clusters | No | No |

*The individual was given these drugs at the hospital before death.
