## Supplementary material for "Spatial multi-omics implicates vascular adipocytes proximal to tertiary lymphoid structures in human aortic aneurysm and B-cell autoreactivity": Table S6

### Human IO Protein Panel Modules

nanoString — GeoMx® TAP Human Immuno-Oncology Content v3

| Immune Cell Profiling Panel (Core) | IO Drug Target Panel | Immune Activation Status Panel | Immune Cell Typing Panel | Cell Death Panel (Early Access) | MAPK Signaling Panel (Early Access) | PI3K/AKT Signaling Panel (Early Access) |
|---|---|---|---|---|---|---|
| Beta-2-microglobulin | 4-1BB CD137 | CD25 | CD14 | BAD | BRAF | p-AKT (S473) |
| CD3 | ARG1 | CD27 | CD34 | BCL6 | p-CRAF (S338) | p-GSK3 (S9) |
| CD4 | B7-H3 | CD40 | CD45RO | BCLXL | EGFR | p-GSK3αβ (S21/S9) |
| CD8 | GITR | CD44 | CD66b | BIM | p-JNK (T183/Y185) | INPP4B |
| CD11c | IDO1 | CD80 | CD163 | CD95/Fas | p-MEK1 (S217/S221) | MET |
| CD20 | LAG3 | CD127 | FAPalpha | Cleaved Caspase 9 | p-P38 (T180/Y182) | Pan-AKT |
| CD45 | OX40L | ICOS | FOXP3* | GZMA | p44/42 MAPK ERK1/2 | PLCG1 |
| CD56 | STING | PD-L2 | | NF1 | p-p44/42 MAPK ERK1/2 (T202/Y204) | p-PRAS40 (T246) |
| CD68 | Tim-3 | | | p53 | p-P90RSK (T359/S363) | p-Tuberin (T1462) |
| CTLA4 | VISTA | | | PARP | Pan-RAS | |
| Fibronectin | | | | | | |
| GZMB† | | | | | | |
| HLA-DR | | | | | | |
| Ki-67 | | | | | | |
| Pan-cytokeratin | | | | | | |
| PD-1 | | | | | | |
| PD-L1 | | | | | | |
| SMA | | | | | | |
| Ms IgG1 | | | | | | |
| Ms IgG2a | | | | | | |
| Rb IgG | | | | | | |
| GAPDH | | | | | | |
| Histone H3 | | | | | | |
| S6 | | | | | | |
